# A reversible memory switch for plant synthetic biology based on the phage PhiC31 integration system

**DOI:** 10.1101/656223

**Authors:** Bernabé-Orts Joan Miquel, Quijano-Rubio Alfredo, Mancheño-Bonillo Javier, Moles-Casas Victor, Selma Sara, Granell Antonio, Orzaez Diego

## Abstract

Plant synthetic biology aims to contribute to global food security by engineering plants with new or improved functionalities. In recent years, synthetic biology has rapidly advanced from the setup of basic genetic devices to the design of increasingly complex gene circuits able to provide organisms with novel functions. While many bacterial, fungal and mammalian unicellular chassis have been extensively engineered, this progress has been delayed in plants due to their complex multicellular nature and the lack of reliable DNA devices that allow an accurate design of more sophisticated biological circuits. Among these basic devices, gene switches are crucial to deploying new layers of regulation into the engineered organisms. Of special interest are bistable genetic toggle switches, which allow a living organism to exist in two alternative states and switch between them with a minimal metabolic burden. Naturally occurring toggle switches control important decision-making processes such as cell fate and developmental events. We sought to engineer whole plants with an orthogonal genetic toggle switch to be able to regulate artificial functions with minimal interference with their natural pathways. Here we report a bistable toggle memory switch for whole plants based on the phage PhiC31 serine integrase and its cognate recombination directionality factor (RDF). This genetic device was designed to control the transcription of two genes of interest by inversion of a central DNA regulatory element. Each state of the device is defined by one transcriptionally active gene of interest, while the other one remains inactive. The state of the switch can be reversibly modified by the action of the recombination actuators, which were administered externally (e.g. via agroinfiltration), or produced internally in response to an inducible chemical stimulus. We extensively characterized the kinetics, memory, and reversibility of this genetic switch in *Nicotiana benthamiana* through transient and stable transformation experiments using transgenic plants and hairy roots. Finally, we coupled the integrase expression to an estradiol-inducible promoter as a proof of principle of inducible activation of the switch.

## MATERIAL AND METHODS

### Construction and assembly of the GoldenBraid phytobricks

A complete list of all phytobricks used in this work can be found in Table S1 and their sequence information and assembly history is available at the GoldenBraid online repository (https://gbcloning.upv.es/). The list of oligonucleotides used to build them can be found on Table S2. The level 0 parts of the PhiC31 integrase and its recombination directionality factor gp3 (RDF) were assembled as previously described to obtain the SET and RESET actuators (GB1531 and GB1531+GB1508 respectively) (1). The registers were designed and assembled as described in Figure S1. In detail, register PB Level 0 part (GB1495) was assembled in a 2 step PCR, (i) addition of attP to the 5’ inverse sequence of TMtb using primers ALF15EN04 and ALF15NOV03 (ii) addition of the attB to the 3’ of P35s (excluding its RBS) using primers ALF15EN05 and ALF15NOV04. Both parts (i) and (ii) were assembled at Level 0 in pUPD2 through a BsmbI-mediated restriction-ligation reaction. Register RL (GB 1514) was amplified from genomic DNA (gDNA) of transiently transformed *N. benthamiana* with register PB and PhiC31 integrase using primers ALF15DIC06 and ALF15DIC07. The reporter genes to be assembled into the register module were amplified from a pre-assembled TU of the GoldenBraid collection as described in Fig. S1B-C. In brief, TUs containing a P35S, followed by a CDS (DsRED, YFP or Fluc) and a Terminator were amplified using primers to obtain a 5’UTR(RBS):CDS:Terminator amplicon to be assembled as a Level 0 part; either to be used as the reverse CDS or direct CDS for the register. Reverse CDSs were amplified using primers ALF15DIC08 and ALF15DIC10 depending on the terminator used in the TU, Tnos and T35s respectively. Direct CDSs were amplified using the primer pairs ALF15DIC09 and ALF15DIC10 depending on the terminator used in the TU, Tnos and T35s respectively. The Register Modules (RMs) were assembled in a Level 1 reaction using three Level 0 parts: (i) reverse CDS, (ii) register PB or RL and (iii) direct CDS. All assemblies were performed as regular GoldenBraid reactions (2).

### Time-dependent characterization of reporter expression in transiently expressed and integrated register modules

Transient expression experiments were performed to test the SET and RESET recombination processes as described in the main text. For the initial characterization by transient expression, equal volumes of *Agrobacterium* cultures transformed with the register modules (GB1523, GB1524, GB1527, GB1528) and their appropriate operators were mixed (GB1531 (SET), GB1508+GB1531 (RESET) or GB0106 (SF negative control) and agroinfiltrated in wild type (WT) *N. benthamiana* plants. For the transgenic lines, only the aforementioned operators or GB0108 (P19) as negative control were used. These cultures were first grown from glycerol stocks for two days until saturation, then 10 uL were sub-cultivated for 16h. Next, the cultures were pelleted, resuspended in the agroinfiltration buffer (10 mM MES, pH 5.6, 10 mM MgCl2, and 200 μM acetosyringone) and adjusted to the appropriated optical density (Table S3). Four to five weeks-old grown *N. benthamiana* plants in a stable condition of 24 °C (light)/20°C (darkness) with a 16-h-light/8-h-dark photoperiod were used. One plant per experimental point was used (1 to 7 days post infiltration (dpi)). Three consecutive leaves (third to fifth from the base) were infiltrated, half with the recombination mixture and half with the negative control. At each time-point, one sample per infiltrated leave was collected using a 0.8 cm cork-borer (approximately 20 mg of tissue for the luminescence quantification assays and 150 mg for the gDNA extraction) and frozen in liquid nitrogen immediately. Samples were grinded with a Retsch Mixer Mill MM400 for 1 min at 30 Hz and stored at −80ºC for subsequent analysis.

### Confocal laser microscopy

Agroinfiltrated leaves of *N. benthamiana* were examined at 5 dpi by confocal laser microscopy with a ZEISS 780 AxioObserver Z1 to visualize the yellow fluorescent protein YFP (λex = 514 nm; λem = 518-562 nm) and DsRED (λex = 561 nm; λem = 563-642 nm) fluorescence. Images of 9-16 tiles were taken to visualize a larger area and processed with ZEN lite 2.5 lite and FIJI softwares.

### Firefly and renilla luciferase luminescence quantification

Firefly lucirerase (Fluc) and renilla luciferase (Rluc) activities were determined using the DualGlo^®^ Luciferase Assay System (Promega) manufacturer’s protocol with minor modifications. Homogenized leaf disc samples were extracted with 375µl of ‘Passive Lysis Buffer,’ vortexed and then centrifuged for 10 min (14,000×g) at 4 °C. Then, 7.5 µl of supernatant were transferred to a 96-well plate and 30 µl of LARII added to quantify the Fluc activity. Finally, 30 µl of Stop&Glow Reagent were used to measure the Rluc activity. Measurements were made using a GloMax 96 Microplate Luminometer (Promega) with a 2s delay and a 10s measurement. Fluc/Rluc ratios were determined as the mean value of three biological replicates coming from three independent agroinfiltrated leaves of the same plant.

### Generation of *N. benthamiana* transgenic plants

Fully expanded leaves were sterilized and used to obtain 0.5 cm diameter leaf-discs with a cork-borer. After an overnight incubation in co-culture medium (MS supplemented with vitamins (Duchefa), 3% sucrose (Sigma-Aldrich), 0.8% Phytoagar (Duchefa), 1 µg/mL BAP (Sigma-Aldrich), 0.1 µg/mL NAA (Sigma-Aldrich)) the leaf-discs were inoculated for 15 min with the *A. tumefaciens* strain LBA4404 carrying the register module construct (Fig. S2, Table S1). Then, the discs were returned to the co-culture medium and incubated for 2 days in darkness. Next, discs were placed in the selection medium (MS supplemented with vitamins (Duchefa), 3% sucrose (Sigma-Aldrich), 0.8% Phytoagar (Duchefa), 1 µg/mL BAP (Sigma-Aldrich), 0.1 µg/mL NAA (Sigma-Aldrich), 500 µg/mL carbenicillin, 100 µg/mL kanamycin. Discs were transferred to fresh medium periodically until the callus and then the explants were formed (5-8 weeks). Explants were cut off and planted in rooting medium (1/2 MS supplemented with vitamins (Duchefa), 3% sucrose (Sigma-Aldrich), 0.8% Phytoagar (Duchefa), 0.1 µg/mL NAA (Sigma-Aldrich), 100 µg/mL kanamycin) until enrooted. All in vitro steps were carried out in a long day growth chamber (16 h light/ 8 h dark, 25ºC, 60-70% humidity, 250 µmol⋅m-2⋅s-1 photons). Samples for phenotyping were collected once the plants were sufficiently developed to harvest 20 mg of tissue to perform the Fluc and YFP quantifications. T1 seeds of Fluc and YFP positive plants were selected for a segregation analysis of the transgene in kanamycin plates. Those scored as a single copy by the chi-square test were used for subsequent experiments.

### Generation of *N. benthamiana* transgenic hairy roots

Wild type and PB LUC:YFP T1-2 *N. benthamiana* plants were used for transformation with *A. rhyzogenes* strain 15834 transformed with the EI LUC or EI PhiC31 constructs (Fig. 5A, Table S1). Fully expanded leaves were used to obtain sterilized leaf-discs which were cultivated for one day in co-culture medium without hormones. Saturated agrobacterium cultures were sub-cultivated and grown overnight. The culture medium was removed by centrifugation at 4.500 rpm for 15 min and then the bacterial pellet resuspended and adjusted with MS to an OD600 = 0.3. Those cultures were employed for inoculation of the sterilized leaf-discs for 30 min under agitation. Next, the leaf-discs were blotted, placed in the co-culture medium and chilled in darkness for 2 days. After that, the discs were transferred to the selection medium without kanamycin nor hormones. The medium was renewed periodically until roots emerged. Transformed roots were selected using a Leica MacroFluo MZZ16F with a DsRED filter. See “Generation of *N. benthamiana* transgenic plants” for the culture medium and growth conditions.

### Estradiol induction experiments

Individual DsRED(+) hairy roots transformed with the EI LUC and EI PhiC31 constructions were divided in two portions and separated in MS medium supplemented or not with 20 μM β-estradiol (Sigma-Aldrich). After 3 days, bright field and red fluorescence images were acquired using a FujiFilm LAS-3000 imager. Then, 100 μM luciferin diluted in MS was added and incubated for 30 min previous acquisition of the chemiluminescence images with the same device. Finally, all the roots were transferred to new MS medium without estradiol and grown for 7 days to repeat the imaging process. For the chemiluminescent imaging the “Ultra” mode was employed with 1 sec of exposition time. Samples of induced roots were collected and stored at −80ºC to perform Fluc/Rluc quantification and PCR-analysis. The grow conditions of “Generation of *N. benthamiana* transgenic plants” applies for this section too.

### YFP quantification

Samples of T0 *N. benthamiana* transgenic plants were used for the YFP quantification using a VICTOR X5 2030 (PerkinElmer), excitation filter F485 and emission filter F535-40. Grinded samples were added with 150 μL of PBS 1X, vortexed and then centrifuged at 13.000 rpm for 10 min at a 4ºC. Finally, 10 μL of the extract were pipetted to a 96-well plate and measured with the spectrofluorometer. The YFP values are expressed as the mean of the measure of three independent leaves from the same plant.

### Genomic DNA extraction and PCR amplification

Genomic DNA was extracted following the C-TAB protocol (3). The obtained gDNA was used for amplification of the register module using MyTaqTM DNA Polymerase (Bioline) and a pair of specific primers for the PB or RL configuration and the PhiC31 recombinase (Table S2). The amplification was confirmed on 1% agarose gel electrophoresis.

## INTRODUCTION

Plant synthetic biology (PSB) is an emerging field that combines the engineering principles of decoupling, abstraction and standardization with plant biology to provide plant systems with new traits and functions (4). Plants naturally offer a myriad of metabolic resources that could be harnessed by modern synthetic biology to tackle current challenges in alimentation, energy production, pest management, or production of valuable biologicals for medicine and industry (5). Recent efforts have been focused on introducing metabolic pathways to create new traits such as vitamin production or glycolate metabolism (6,7), fixation of atmospheric nitrogen (8), increased yield or improving photosynthesis (9,10). Nevertheless, the implementation of these new functions requires precise regulation of the new genetic networks to ensure an optimal use of the plant resources and avoid deleterious effects. The introduction of complex programmed genetic networks in bacterial and mammalian systems has been a great success, achieving engineered organisms that behave as cellular computing devices that have substantial biotechnological and therapeutic impact (11). This success has been possible due to the previous development and extensive characterization of genetic devices and molecular tools that, combined, generate increasingly complex functions. However, the translation of these complex genetic circuits to plants has been challenging due to the limited availability of well-documented genetic elements as well as the technical limitations of high-throughput plant transformation and circuit characterization (4,12). Current efforts in PSB are focused on developing reliable DNA assembly standards (phytobricks) to allow more efficient biodesign and characterization of plant functions (1,13).

Among the diverse synthetic biology tools developed in the past years, basic memory devices as e.g. transcriptional toggle switches have been pivotal in allowing the design of increasingly complex genetic circuits (14). Traditional inducible control systems lacking memory functions require a constant supply of actuators (activators or repressors) for a sustained control of cell outputs, often imposing a metabolic burden to the cell (15). In contrast, toggle switches provide the ability to respond to punctual external inputs with a sustained response. Applied to crop plants, gene memory devices could facilitate the deploy of new layers of genetic control over agronomically-relevant outputs such as flowering time, stress response or the biosynthesis of added value metabolites. Furthermore, memory systems circumvent the need for continuous addition and monitoring of external inducers, which can be expensive and difficult to control, especially in large settings as crop fields. While several non-memory chemically-inducible systems have been adapted to plant biotechnology, such as ethanol (16), glucocorticoids (17,18), copper (19,20) or insecticides as methoxyfenozide (21), memory devices adapted and/or engineered for the plant chassis are almost absent. As a notable exception, Müller *et al*., 2014 (22) developed a red light-inducible toggle switch for plant cell protoplasts, based on the interaction of *Arabidopsis thaliana* PhyB-PIF6 proteins. This approach showed remarkable spatiotemporal resolution of gene expression. However, this system relies on light for activation, thus, its adaptation to whole plants can be challenging in applications in which the illumination conditions cannot be strictly controlled, as in open field applications. Additionally, the memory of this optogenetic system is based on the slow dissociation rate of the PhyB-PIF6 protein-protein interaction, which will result in eventual inactivation due to the decay and dilution of the protein components, making this system unsuitable for applications in which information needs to be maintained for long periods of time or transmitted over generations. Therefore, alternative gene memory switches for plants that allow long-term and inheritable memory storage are critically needed to enable the advancement of the field.

Synthetic genetic memory can be built through diverse mechanisms, such as transcription-based double negative feedback loops, positive feedback loops or DNA recombination (23,24). Integrases are a group of DNA-actuating enzymes found in temperate bacteriophages that catalyze site-specific recombination (SSR) (25). Serine integrases catalyze the recombination of attP and attB attachment sites, generating hybrid attR and attL sites in a strictly unidirectional reaction. This process can be reversed in the presence of an excisionase or recombination directionality factor (RDF), which in combination with the integrase can catalyze the recombination of the attR-attL sites into attP-attB again (26). This specific DNA recombination event can result in a variety of DNA-rearrangements (integration, excision, inversion and translocation) depending on the topology of the initial DNA molecules and the orientation of the recombination sites. SSR mechanisms of diverse integrases have been extensively exploited in bacteria and mammalian cells to engineer reversible synthetic memory devices and complex logic circuits. Examples include counting cellular events (27), store and rewrite biological data in the chromosome (28), control the flow of the RNA polymerase along the DNA and thus amplifying the expression of a reporter gene (29) and to engineer complex regulatory circuits (30–32). Although SSR systems have been used in plants before, all the efforts have been focused in transgene engineering for crop breeding rather than in the design of memory devices for the control of gene expression (33,34). Some of their uses include the removal of foreign DNA (35), stacking agronomic-valuable traits (36), and chloroplasts engineering (37).

Here, we created the first toggle memory switch for whole plants based of the phage PhiC31 serine integrase and its cognate recombination directionality factor, RDF (38). The switch was designed to control the transcription of two genes of interest, one of which is initially ON, while the second one remains in OFF status. The state of the switch changes by punctual supply of integrase to the cell, and the memory of the new status is maintained until intentionally reversed by a combined new supply of integrase and RDF. The components of the genetic switch were designed for plant expression and standardized by using the GoldenBraid (GB) DNA assembly platform and software to allow easy adaptation of this tool to other PSB applications (1). We extensively tested the kinetics, memory and reversibility of this genetic device in *Nicotiana benthamiana* through transient and stable transformation experiments using transgenic plants and hairy roots. Additionally, we coupled the integrase expression to an estradiol-inducible promoter as a proof of principle of externally-induced activation of the switch.

## RESULTS

### Design of a modular reversible genetic switch for plant systems

To design a reversible genetic switch for plant expression, we adapted a previously reported strategy (28) to the phage PhiC31 integrase and its cognate RDF (38). This genetic switch comprises a flippable DNA promoter element flanked by the PhiC31 attP and attB recombination sites in opposing orientations, named as register PB (Figure 1A). In the initial state, the switchable promoter drives the expression of a gene of interest (GOI2) positioned downstream of the register PB in the direct DNA strand. The orientation of the promoter can be switched by action of the PhiC31 integrase, which catalyzes attPxattB recombination into attR and attL respectively (SET), flipping the DNA segment resulting in a new register RL. As a result, the register RL now drives the expression of a second GOI located in the reverse strand (GOI1), while GOI2 expression has been turned off. Our goal was to design a reversible genetic switch, therefore the register RL can be reversed to the PB state by the combined action of the PhiC31 integrase and RDF factor, which catalyze attRxattL recombination into attP and attB (RESET), resulting in the reconstitution of the initial register PB and expression of GOI2 while GOI1 is inactivated. To maximize the exchangeability and reusability of this this genetic device for PSB applications, we took advantage of the modularity of GB to structure our genetic switch in three standard and interchangeable parts: (i) the direct coding sequence (CDS) encoding GOI2, (ii) the registers PB or RL, and (iii) the reverse CDS encoding GOI1 (Fig. S1A). Reverse and direct CDSes can be easily created from any Transcriptional Unit (TU) conforming the phytobrick standard, as e.g. those available at the GB collection (Fig. S1B and S1C). The register comprised the strong CaMV35S promoter (P35S) and an upstream tomato Metallothionein B terminator (MTB) in opposite orientation as an insulator to avoid any leaky backward expression. Recombination sites were inserted flanking the terminator and promoter elements. Since incorrect insertion of the att site close to the transcription start site (TSS) or ribosome binding site (RBS) could hinder efficient expression, this was inserted in the 5’ untranslated region (5’UTR) of the P35S, downstream of the TATA box and upstream of the RBS, keeping them intact (Fig. S1A). The registers were named according to the 5’-3’ order of the att sites: register PB (attP-attB sequences, GB1494) and the register RL (attR-attL, GB1506). The three elements can be assembled in a single step Golden Gate-like reaction to create the so-called register module (RM). The actuators of the SET and RESET operations were adapted for *in planta* expression and incorporated to the GB collection: Pnos:PhiC31:Tnos (GB1531) and P35S:RDF:T35S (GB1508).

**Figure 1:**
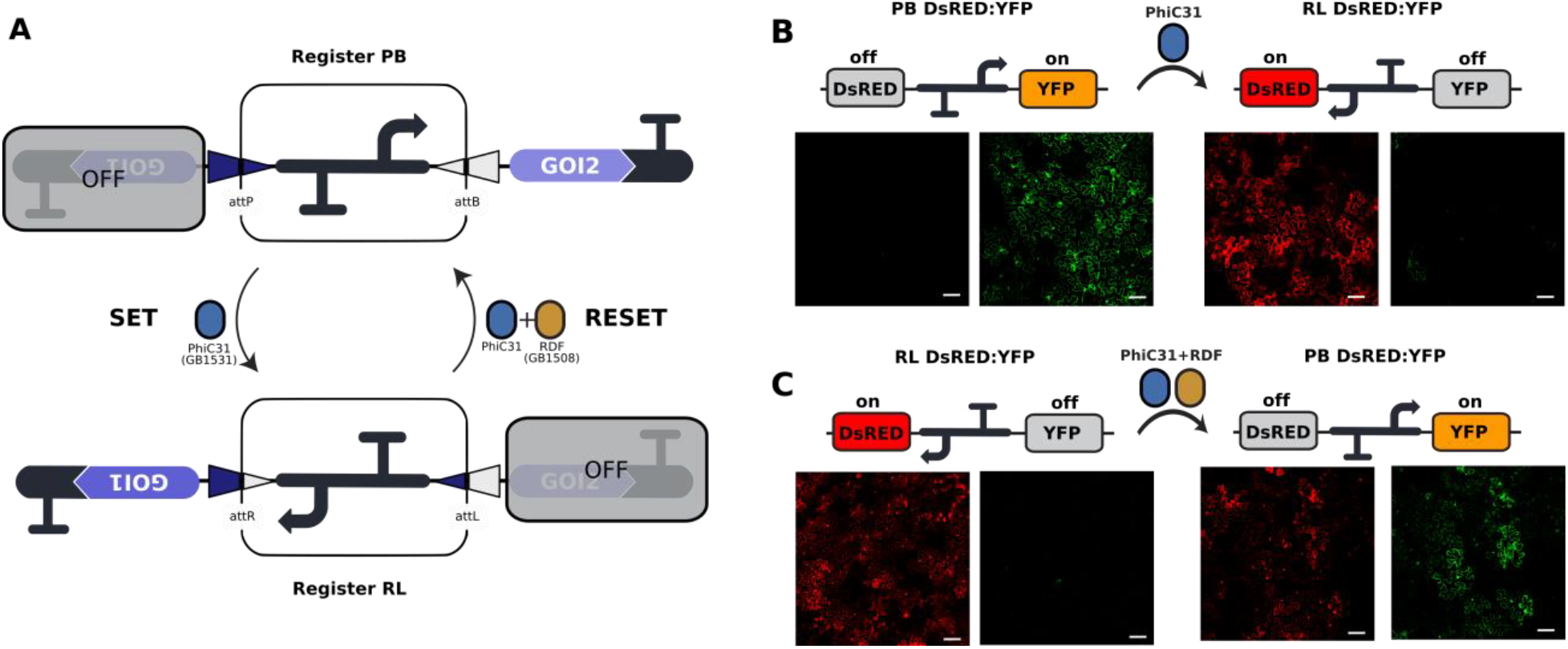
Design and functional validation of the plant toggle switch based on the phage PhiC31 integration system. **(A)** An invertible plant promoter element (Register) works as a toggle switch for the regulated expression of two genes of interest (GOI1 and GOI2). The promoter can be inverted by action of the PhiC31 integrase (SET) which catalyzes site-specific recombination of the attP and attB sites flanking the promoter. This event results in a change in the expression status of the two GOIs and the creation of the chimeric attR and attL sites. Expression of the integrase and recombination directionality factor (RDF) catalyzes recombination of attR and attL to reset the toggle switch to its original state (RESET). The genetic parts encoding the PhiC31 integrase and RDF can be found in the GoldenBraid collection, GB1531 and GB1508 respectively. **(B, C)** CLM images of WT *N. benthamiana* leaves infiltrated with the toggle switch alone (left) or in combination with the recombination actuators (PhiC31 or PhiC31 + RDF), (right). Reporter genes expression (DsRED and YFP) were analyzed in the initial and the commuted states of the switch during the SET (B) and RESET operations (C). Images were taken 3 days post-infiltration. The scale bar represents 100 μm.

To test the functionality of the toggle switch in plants, yellow fluorescent protein (YFP) and DsRED were selected as direct and reverse reporters respectively. The reporters were then combined with the PB and RL registers to create the PB DsRED:YFP and RL DsRED:YFP RMs. Next, we performed *Agrobacterium-*mediated transient expression in *N. benthamiana* leaves by co-infiltrating each RM (PB or RL DsRED:YFP) with its appropriate actuators (PhiC31 or PhiC31+RDF) to evaluate the SET and RESET operations by confocal laser microscopy (CLM), (Fig. 1B-1C). As expected, leaf agroinfiltration of the PB DsRED:YFP register module alone yielded bright YFP expression, with no DsRED expression detected. In contrast, co-infiltration of PB DsRED:YFP with PhiC31 integrase (SET operation) resulted in robust activation of DsRED expression and de-activation of YFP expression in most cells, as shown in Fig. 1B. Similarly, agroinfiltration of RL DsRED:YFP module alone produced only DsRED (+) cells, whereas co-expression of RL DsRED:YFP + Integrase + RDF (RESET operation) resulted in activation of YFP expression (Fig. 1C). As can be observed, the RESET process was not complete, resulting in some cells still expressing DsRED. Co-expression of RL DsRED:YFP with PhiC31 alone did not change the reporter expression, proving the necessity of RDF to enable attRxattL recombination (data not shown). Altogether, with these results we obtained a static picture of the two states of our plant-customized toggle switch, confirming its functionality.

To monitor quantitatively the expression kinetics of the SET and RESET operations, DsRED was substituted by the firefly luciferase (LUC or Fluc) as a reporter gene, producing four new RMs, namely PB LUC:YFP, PB YFP:LUC, RL LUC:YFP and RL YFP:LUC (Fig. 2 and Fig. S2). Additionally, a constitutively-expressed renilla luciferase (Rluc) was assembled to each RM as internal reference (Fig. S2). RMs were transiently co-expressed in WT *N. benthamiana* leaves with its appropriate effector (PhiC31 for SET or PhiC31+RDF for RESET) or with a non-coding DNA fragment (stuffer fragment, SF) as a negative control, and luciferase levels were recorded over time starting at 24 hours post-infiltration (hpi). As shown in Fig. 2A, SET Fluc activation was strong and rapid, producing Fluc/Rluc levels of 43-fold reporter induction from the start of the measurement, indicating that recombination was efficiently taking place almost concomitantly with the transient transformation process. RESET activation of Fluc in the RL YFP:LUC module was also readily detectable at 24 hpi (Fig. 2C), however in this case the maximum Fluc/Rluc signal of 35-fold induction was observed at 60 hpi, indicating slower kinetics of this operation involving two actuators. Fig. 2B and 2D show the deactivation of Fluc using SET and RESET operations respectively. In both cases only partial deactivation of Fluc was observed, reaching approximately 75% signal decrease in the case of SET and only 40% Fluc deactivation for RESET as compared to non-operated agroinfiltrated control.

**Figure 2:**
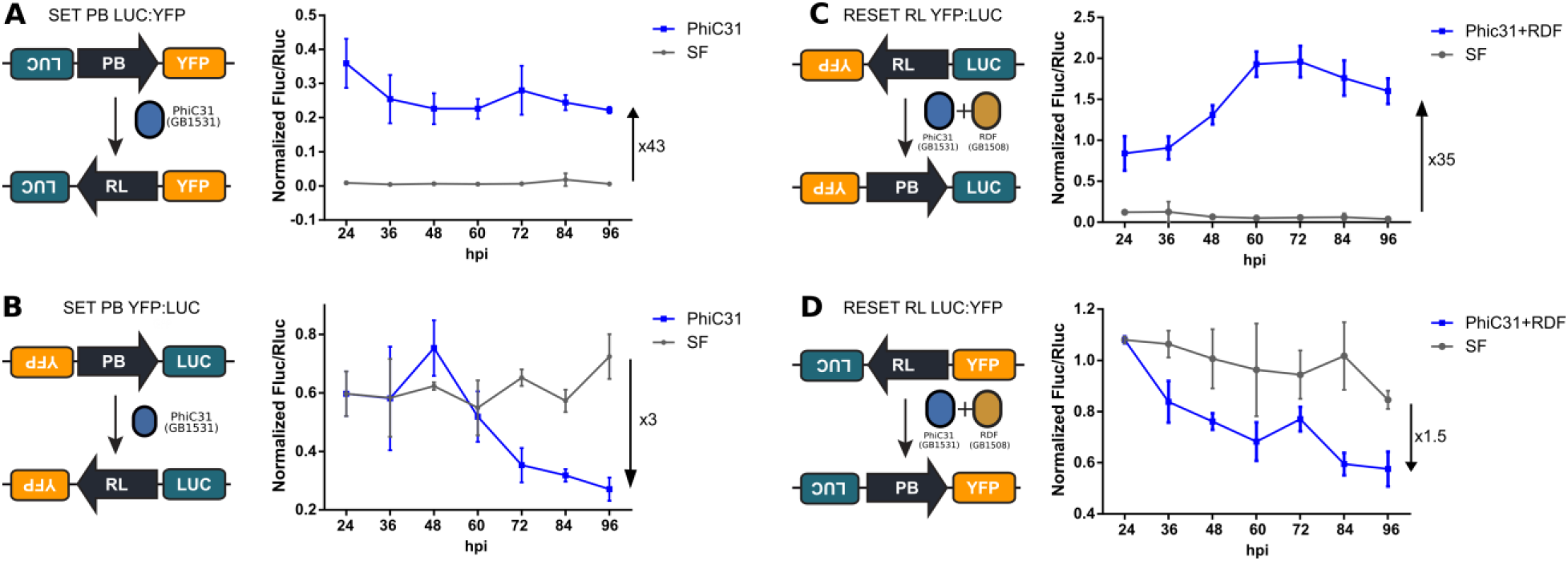
Evaluation of the SET and RESET operations on RMs by transient expression in WT *N. benthamiana* leaves. **(A-B)** SET of the PB LUC:YFP (A) and the PB YFP:LUC (B) mediated by the PhiC31 integrase. **(C-D)** RESET of the RL YFP:LUC (C) and RL LUC:YFP (D) mediated by the combined action of PhiC31 integrase and RDF. **(A, C)** The recombination processes switch on Fluc reporter expression resulting in an increase of the Fluc/Rluc ratios (Rluc was expressed as a constitutive internal control). **(B, D)** The recombination reactions switch off Fluc expression resulting in a decrease of the Fluc activity over the time. All the graphs represent the evolution of Fluc/Rluc ratios at intervals of 12 hours post infiltration (hpi) for 96h. Leaves were treated with the recombination actuators (blue line) or with an empty vector named SF used as negative control (grey line). Each point represents the mean of Fluc/Rluc ± SD of three leaves in different plants.

### Characterization of the register modules in stably transformed *N. benthamiana* plants

While transient transformation enables quick characterization of molecular tools developed for plants, we sought to develop a genetic switch that could be stably introduced into genetically modified plants to suit multiple applications. *Agrobacterium*- mediated stable transformations involve the random integration of foreign transfer DNA (T-DNA) in the plant genome, which may result in unpredicted changes in the performance of the genetic part. In order to further characterize the performance of our genetic switch in the genomic context, we generated transgenic *N. benthamiana* lines carrying the four aforementioned RMs (PB LUC:YFP, PB YFP:LUC, RL LUC:YFP and RL YFP:LUC; Fig. S2). T0 plants were screened based on YFP and Fluc expression levels, and then one line of each RM with a single copy of the transgene was selected to perform subsequent experiments on its T1 progeny (Fig. S3). SET and RESET operations were performed by *Agrobacterium*-mediated transient delivery of the recombination actuators. Optimal levels of each actuator were experimentally determined by titrating the optical density (OD) of the respective *Agrobacterium* cultures, a parameter which is known to correlate with the amount of T-DNA delivered to the transformed cells (1) (Fig. S4). Agroinfiltrated leaves were sampled and analyzed for Fluc and Rluc expression every 24h for 7 days (Fig. 3). As anticipated, the RMs where Fluc expression turns on upon performing operations (SET PB LUC:YFP and RESET RL YFP:LUC) responded to actuator delivery with a rapid increase in Fluc/Rluc levels, reaching a maximum at 5 days post infiltration (dpi). Concomitantly with Fluc activation, YFP levels decreased markedly when measured in the same infiltrated areas at 5 dpi (Fig. 3A and 3C, Fig. S5B). Reporter Fluc/Rluc induction levels for the transgenic lines were 13-fold and 15-fold for the SET and RESET operation respectively. Additionally, while Fluc levels were sustained over time for the SET operation (Fig. 3A), Fluc/Rluc signal showed a slow decrease after 5 dpi for the RESET operated RMs (Fig. 3C). This could be caused by an excess of PhiC31 integrase not bound to RDF, which would cause multiple RESET/SET recombination cycles decreasing the stability of the measurements. On the other side of the equation, those RM conformations where Fluc turns off upon commutation (SET PB YFP:LUC and RESET RL LUC:YFP), showed a moderate but consistent decrease of Fluc/Rluc levels in parallel to a clear activation of the YFP signal when commuted by agroinfiltration (Fig. 3B and 3D, Fig. S5A). Altogether, these results show that the SET or RESET operations take place correctly in the genomic context of stably transformed plants.

**Figure 3:**
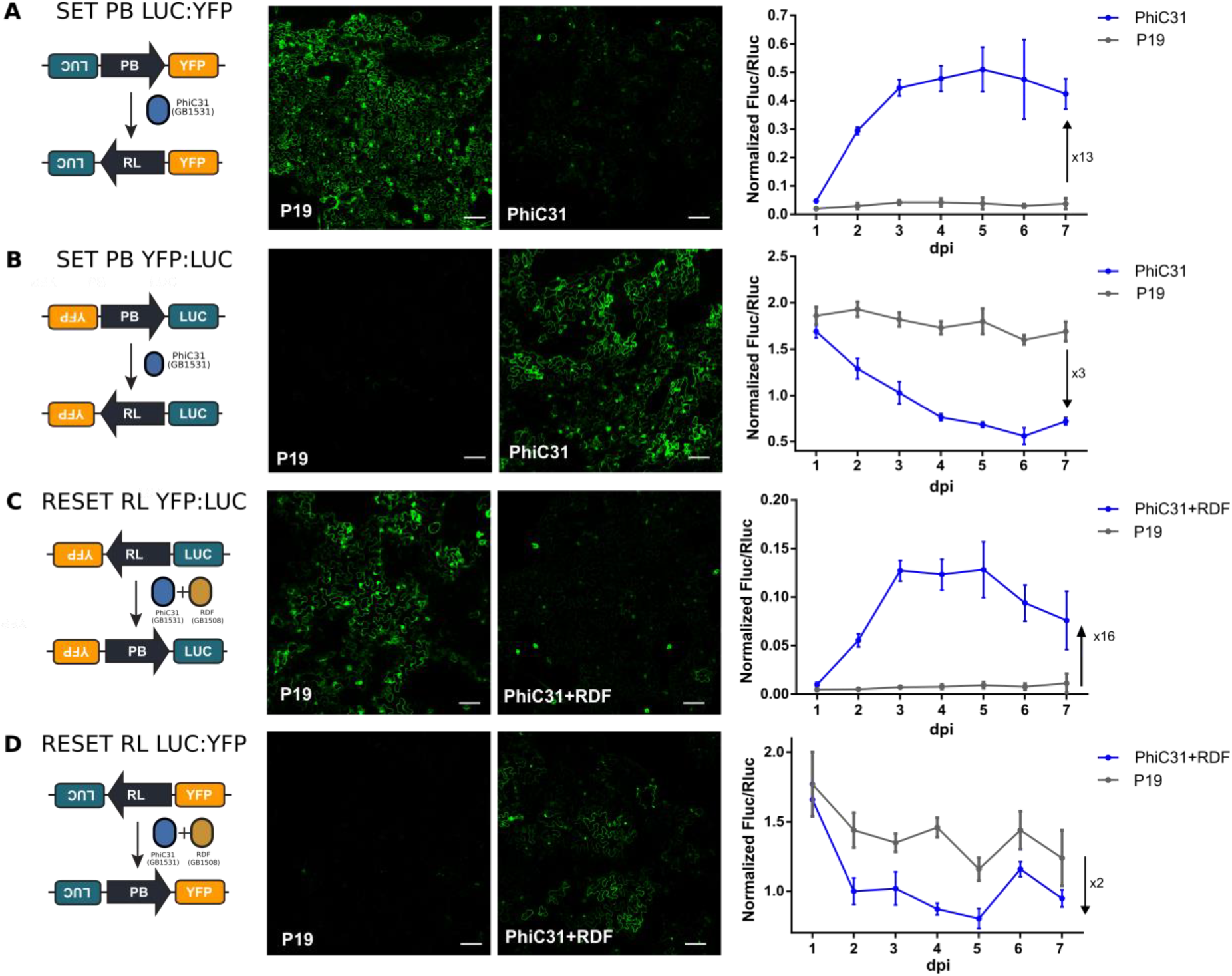
Evaluation of the SET and RESET operations in transgenic *N. benthamiana* with RMs integrated in their genome. **(A-B)** SET recombination of the PB LUC:YFP (A) and PB YFP:LUC (B) mediated by the PhiC31 integrase. **(C-D)** RESET recombination process mediated by the combined action of PhiC31 and RDF of the RL YFP:LUC (C) and RL LUC:YFP (D) RMs. **(A,C)** In the initial state the YFP reporter is actively expressed as seen by the CLM image infiltrated with P19 (negative control). Infiltration of the recombination actuators flips the register cassette inactivating YFP expression (PhiC31 or PhiC31+RDF images) and activating the Fluc expression (graph, blue line). **(B, D)** In the initial state YFP expression is off as demonstrated by the P19 control. Agroinfiltration of actuators switches on YFP expression (PhiC31 or PhiC31+RDF images) and switches off Fluc expression (graph, blue line). CLM images were taken at 5 days post infiltration (dpi); scale bars represent 100 μm. All the graphs represent the evolution of Fluc/Rluc values for 7 dpi of leaves transiently transformed with actuators (blue lines) or P19 as negative control (gray line). Experimental points show the mean of normalized Fluc/Rluc values of three agroinfiltrated leaves ± SD.

### Stable and reversible memory storage over a full SET/RESET cycle in whole plants

A distinctive feature of a PhiC31-based memory device is the long-term stability of commutations based on DNA recombination (39). Our primary goal was to create a switch that could alternate between two different configurations and whose memory status could be transmitted to the progeny. To this end, we set out to test the memory and reversibility of the switch by performing a full SET/RESET cycle in *N. benthamiana* lines with the RMs integrated in the genome. As successive agroinfiltrations of the actuators would result in excessive damage to the plants for accurate analysis, we decided to generate commutated calluses after the first SET operation, to then perform the second RESET operation on fully regenerated plants. In the first place, the efficiency of the SET and RESET operations was tested in the context of *in vitro Agrobacterium*-mediated plant leaf disc transformation. RL LUC:YFP and PB YFP:LUC lines were used to follow the course of the transformation through YFP expression activation (Fig. S6A). This approximation showed that the SET operation produced twice as much fluorescent callus than the RESET (Fig. S6B-C). However, these transformations ended with highly chimeric calluses which eventually would yield plants with mixed states of the RMs, probably because the recombination took place in an advanced stage of the callus generation process. We reasoned that performing the SET or RESET operation by agroinfiltration prior to the *in vitro* regeneration would increase the initial population of commutated cells, raising the chances to obtain fully commutated calluses and subsequently fully commutated plants. For the same reason, we limited the experiment to the SET operation which showed to be more efficient. Therefore we conducted the experiment illustrated in the Fig. 4A, using the aforementioned PB YFP:LUC stable line to perform a SET operation by transient expression of PhiC31 in the leaf. Then, at 5 dpi when the peak of recombination was reached, fluorescent discs were excised and regenerated *in vitro* to obtain stably switched calli. As previously shown, this resulted in chimeric calli with variable efficiency of the SET operation and thus, variable YFP expression levels. Then, the calli were differentiated to obtain explants, which resulted in seven regenerating plant lines (P1-P7), one of which (P3) died during the in vitro cultivation process. In order to assess the configuration of the RM in each line (either the initial PB YFP:LUC or the commutated RL YFP:LUC state), regenerating lines were genotyped using specific primer pairs for PB or RL state. The presence of integrated PhiC31 was also analyzed in the same lines (Fig. 4B). Plants P1 and P2 did not show evidence of commutation events (only PB configuration detected). P1 showed no presence of PhiC31, while P2 showed only a weak band. P5 and P6 were apparently chimeric plants in which both configurations coexisted, P6 showing only a faint band for commuted configuration compatible with the absence of PhiC31 integration. Finally, P4 and P7 were positive only for RL and PhiC31, indicating a complete switch to the RL configuration. Thus, P7 explant was transferred to soil and used to test the reversibility of the genetic device.

**Figure 4:**
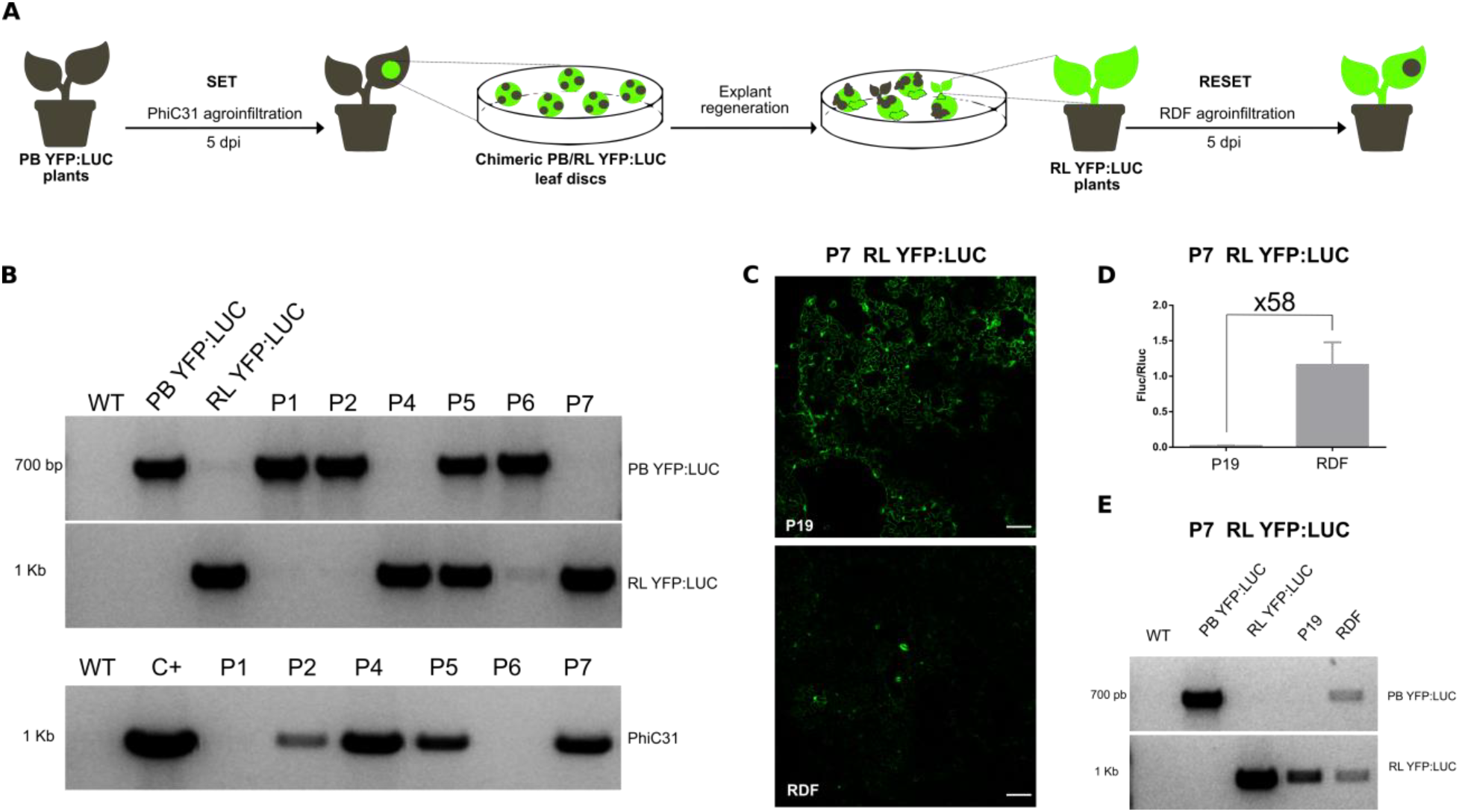
Full SET/RESET cycle of the plant toggle switch over a plant transformation and regeneration. **(A)** Representation of the experiment conducted to assess the bistability and reversibility of the toggle switch in *N. benthamiana* PB YFP:LUC transgenic lines (YFP off, Fluc on). PB YFP:LUC plants were agroinfiltrated with the PhiC31 integrase to induce a SET to RL YFP:LUC. Five days post infiltration fluorescent leaf discs were collected, sterilized and then cultivated *in vitro* until explant regeneration. Explants genotype was then analyzed by PCR, to select totally commutated RL YFP:LUC plants. RL YFP:LUC plants (such as P7) were then agroinfiltrated with PhiC31+RDF to operate the RESET to the original PB YFP:LUC configuration and demonstrate the reversibility of the system. **(B)** Agarose electrophoresis gels showing the genotyping results of the regenerated plants (P1-P7). The upper panel reflects the results for a specific pair of primers for the amplification of the original PB YFP:LUC configuration. The middle panel shows the results when amplifying with a specific pair of primers for the commutated RL YFP:LUC state. The lower panel shows amplification results of integrated PhiC31 integrase. **(C)** CLM images of agroinfiltrated P7 with P19 (negative control) and RDF (RESET) after 5 dpi. Scale bars represent 100 μm. **(D)** Fluc/Rluc ratio of P7 agroinfiltrated with P19 or RDF at 5 days post induction. Values represent the mean Fluc/Rluc values for three different agroinfiltrated leaves ± SD. **(E)** Genotyping results of agroinfiltrated P7 with P19 and RDF at 5 dpi.

At maturity, P7 leaves were agroinfiltrated with RESET effector (RDF) or with a control (P19) and analyzed for YFP (Fig. 4C) and Fluc expression (Fig. 4D). Remarkably, YFP expression significantly decreased at 5 dpi in RESET samples (Fig. 4C) and Fluc activity was induced 58-fold over the un-induced background levels, reaching approximately 60% of the Fluc/Rluc ratios typically observed in the original PB YFP:LUC plant (Fig. 4D). These results were also genotypically confirmed by PCR-amplification of the commuted tissue, as compared to control samples (Fig. 4E). Altogether, these experiments showed that the toggle switch, once integrated in the genome can effectively perform a full cycle of SET/RESET operations.

### Chemical induction of PhiC31 controls SET operation in *N. benthamiana* hairy roots

Master regulators are essential for engineering synthetic gene networks in which metabolic fluxes can be deviated among different pathways. So far, we have demonstrated that our plant-custom-made PhiC31-based toggle switch can alternate between two defined states and maintain memory, a key feature of a master regulator. However, we relied on agroinfiltration to induce a state change by expression of the SET and RESET effectors, which does not allow precise spatiotemporal control of expression. We sought to achieve tighter control over the recombination process by coupling the expression of PhiC31 to an estradiol-inducible system, which was named estradiol-inducible PhiC31 (EI PhiC31). To this end, a GB module was assembled, encompassing three transcriptional units encoding for (i) a chimeric trans-activator (estradiol receptor (ER) fused in N-terminal to the LexA binding domain (LexABD) and to GAL4 activation domain (GAL4AD) in C-terminal); (ii) an inducible PhiC31 with the LexA operator and the mini35S promoter; and (iii) a DsRED fluorescent marker (Fig. 5A). In this system, the trans-activator is constituvely expressed and remains in the cytoplasm. Upon addition of estradiol, the trans-activator localizes to the nucleus and binds to the DNA operator, inducing the expression of PhiC31 integrase. Expression of PhiC31 results in a SET operation which is evidenced by Fluc luminescence.

**Figure 5:**
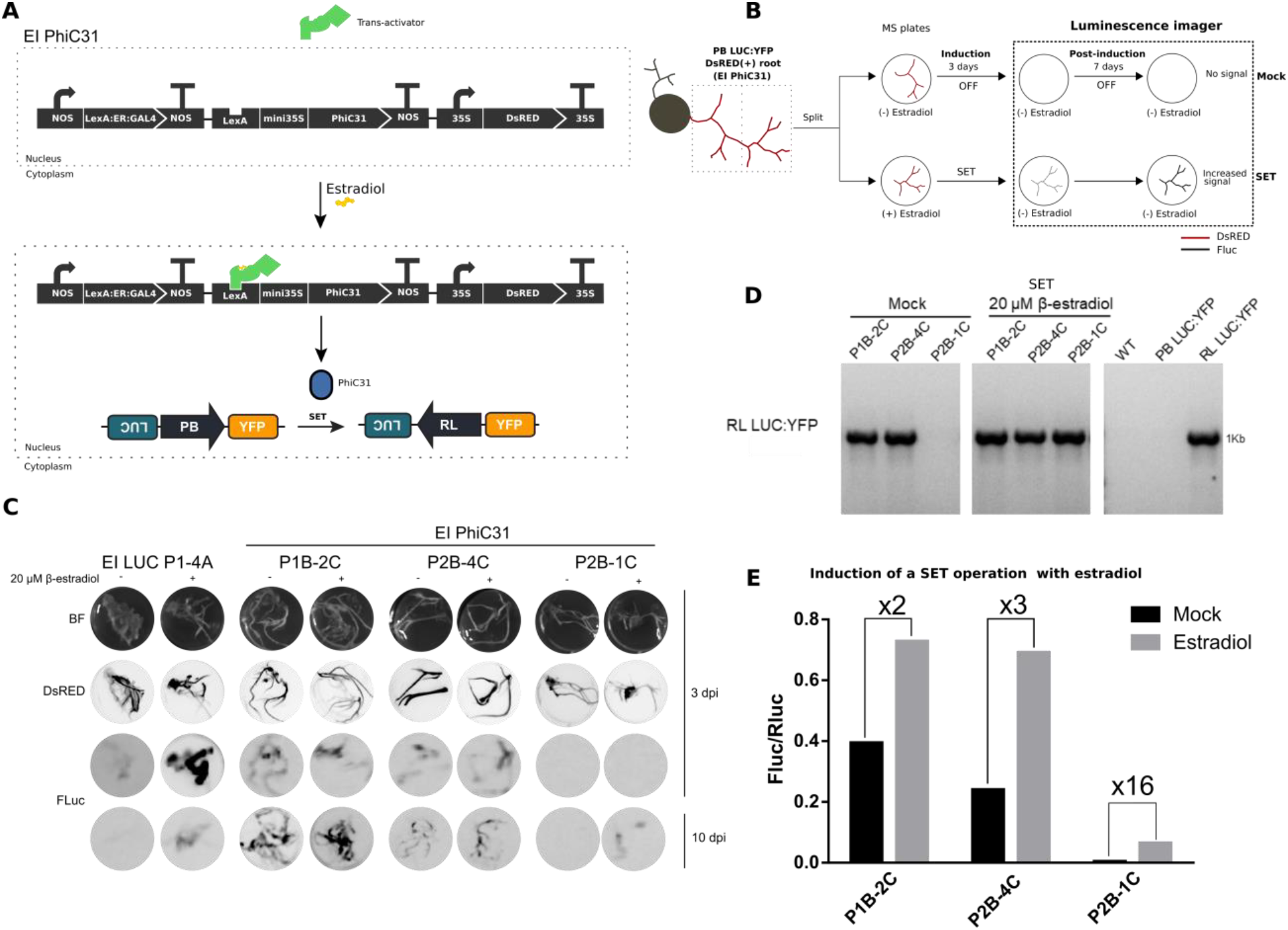
Chemical induction of the PhiC31 recombinase in stable *N. benthamiana* PB LUC:YFP hairy roots. **(A)** Representation of the estradiol-inducible (EI) system EI PhiC31. In absence of estradiol, the constitutively expressed chimeric trans-activator is confined to the cytoplasm. Upon addition of estradiol it localizes to the nucleus where it induces the expression of the PhiC31 integrase. This enables the SET operation of the PB LUC:YFP register module turning on Fluc expression. **(B)** Diagram of the EI PhiC31 experiment in hairy roots. DsRED(+) *N. benthamiana* PB LUC:YFP hairy roots expressing EI PhiC31 were divided and incubated in MS plates in presence (SET) or absence of estradiol (Mock) for 3 days. After chemoluminescence imaging, roots were transferred to new estradiol-free plates where they remained for 7 days before imaging them again to measure Fluc activity. **(C)** Images of the EI LUC and different EI PhiC31 hairy roots used in the induction experiment. First row images correspond to bright field images (BF), second row are fluorescence images to detect the DsRED marker, third and fourth are luminescence images (FLuc). A LAS3000 imager was used to take all the images at 3 and 10 dpi. Chemoluminescence images were taken with the “Ultra” mode and 1 sec of exposition. **(D)** Genotyping results of uninduced (left panel) and induced (central panel) EI PhiC31 hairy roots, including negative (WT and PB LUC:YFP) and positive (RL LUC:YFP) controls (right panel). A specific pair of primers for RL LUC:YFP amplification was used. **(E)** Quantification of the Fluc/Rluc ratios of induced and mock EI PhiC31 hairy root lines. Bars represent the mean of three technical replicates for an individual root.

We used *A. rhyzogenes* to stably transform the EI PhiC31 system into the previously described PB LUC:YFP line. This transformation system enables the regeneration of roots which are an ideal system for high-throughput chemical screening. We performed the estradiol-induction (EI) experiments as shown in Fig. 5B. Successfully transformed DsRED(+) roots were incubated with or without estradiol. The roots were then imaged for luminescence resulting from activity of the Fluc reporter. Subsequently, the roots were transferred to new estradiol-free MS plates and incubated for 7 days before imaging again. In parallel, WT *N. benthamiana* was transformed with a construct bearing an estradiol-inducible Fluc reporter as a positive control with no memory device (EI LUC). As shown in Fig. 5C and Fig. S7, roots transformed with EI LUC had a quick increase of luminescence at 3 days post-induction (3 dpi) and lost most of the signal at 10 dpi. On the other hand, EI PhiC31 root lines maintained activity over time up to day 10, as expected for a memory switch. Unexpectedly, several EI PhiC31 root lines (depicted here lines 2C and 4C) showed high Fluc background in the absence of estradiol, probably due to leaky activation of the estradiol inducible system. Only weak Fluc lines (line 1C) showed no leaky background, and although Fluc intensity at 3dpi was almost undetectable, it reached moderate levels at 10dpi, indicative of a memory-based sustained activation. In order to confirm this, we performed specific PCR-amplification of the RL LUC:YFP configuration (Fig. 5D). P1B-2C and P2B-4C showed RL LUC:YFP amplification before and after induction with estradiol, confirming that the leakiness was caused by an unwanted early switching of the RM. On the other hand, P2B-1C showed no amplification of RL LUC:YFP before induction but strong amplification after induction with estradiol. This result supported our hypothesis that P2B-1C increase in Fluc activity was due to an estradiol-inducible SET operation of the switch. Subsequently, we quantified the luminescence at 10 dpi comparing induced and non-induced roots for each line (Fig. 5E). P1B-2C and P2B-4C lines showed a slight increase of 2-3 fold after induction, while P2B-1C showed a 16-fold induction and a substantially lower Fluc/Rluc ratio compared to the leakiness-induced roots. In conclusion, these results show that a chemically-inducible PhiC31 actuator that can control the activation of a genetic memory switch is achievable. However, further optimization of the induction system is required to avoid leaky expression of the PhiC31 integrase. Factors as the genomic localization of the transgene or its copy number could result in low basal or sporadic expression of the PhiC31 integrase, resulting in an unwanted permanent change in the toggle switch.

## DISCUSSION

Here we sought to expand the molecular toolbox for PSB by designing a bistable and reversible toggle switch for whole plants. Tools to engineer genetic memory in plants are currently limited, being a red-light controlled memory switch for protoplasts the only option available (22). While this system provides great spatiotemporal resolution of gene induction, its adaptation to full plants is challenging due to strict light requirements as well as limitations performing long-term and inheritable memory storage. Serine integrases are powerful tools that can induce a stable change in the DNA and have extensively been used for many applications, including the design of toggle switches for bacteria and mammalian cells (40). We adapted the previously described bacterial switch by Bonnet et al., 2012, which was based on the Bxb1 phage integration system (28). We decided to use the PhiC31 phage integrase and RDF because this integrase has been previously used for genomic engineering in plants, and also suited better the grammar of our GB cloning system. The architecture of the switch was designed to be fully modular, so it can be easily adapted to control the expression of any GB-domesticated downstream gene by performing a single-step GB assembly reaction.

We built multiple RMs to characterize the behavior of the switch at the SET and RESET operations. First, the genetic parts were tested by transient expression of the RMs and the SET and RESET actuators (PhiC31 and PhiC31+RDF), and the evolution of reporter expression in WT *N. benthamiana* was analyzed, both transiently as well as in stably-transformed plants. In all experiments, the initial state of the switch showed strong expression of the active reporter while no noticeable backward expression of the inactive reporter was observed by luminescence or CLM imaging. This suggests that the design of the genetic switch successfully avoided leaky backward expression. When the actuators were applied via agroinfiltration, we observed good performance of reporter activation either by SET or RESET operation. Activation reached up to 43-fold over uninduced controls by transient expression and 15-fold (60% of the fully active state) in the case of transgenic lines, and it was sustained over days. However, when analyzing inactivation of a reporter gene after SET or RESET, we observed slow decrease in the signal, in occasions reaching only a 40-75% decrease over the non-commutated switch. This effect was probably due to the combined effect of slow decay of the reporters and the incomplete switching on the RMs of all the cells analyzed, especially during the RESET operation involving RDF. Phage RDFs have been studied due to their ability to reverse the directionality of a SSR reaction via direct interaction with the cognate integrase, i.e. to catalyze attRxattL reaction (38,41). However, this reaction has shown to be inefficient when the stoichiometry of the integrase and RDF is not optimal, because integrase unbound to the RDF can catalyze the attPxattB reaction, entering a bidirectional cycle of PB-RL-PB recombination. Bonnet et al., 2012 also observed this effect when developing their bacterial toggle switch based on Bxb1 and Xis (RDF) as separate TUs. In fact, these authors postulated that the system enters in a bidirectional regime if the concentration of the excisionase is too low relative to the integrase. Therefore, these authors tested several combinations of genetic constructs to increase the excisionase expression levels and reduce the half-life of the integrase achieving an unidirectional reaction with 90% efficiency. We followed a similar approach by increasing RDF expression levels through the CaMV35 promoter, which has about 10-fold higher expression than the Nopaline Synthase promoter used to express the PhiC31 integrase (42). Additionally, we optimized the OD600 of the *A. tumefaciens* cultures during transient expression to optimize the efficiency of the RESET operation. However, we still saw incomplete attRxattL recombination of all induced cells, as seen by fluorescent imaging of leaf cells and transformed calluses. Recently, Olorunniji et al., 2017 reported higher attRxattL recombination efficiency by creating a fusion of PhiC31 integrase to RDF (43). Using this new Integrase-RDF fusion as the RESET operator could help to solve bidirectionally issues in a future iteration of this plant switch, achieving higher efficiency off attRxattL recombination without the need of high throughput testing of multiple genetic constructs to balance expression levels of both components.

After individual characterization by agroinfiltration of the SET and RESET operations, we decided to test the performance of a full SET/RESET cycle to evaluate the memory of the switch over plant transformation and regeneration. However, a drawback of the agroinfiltration method is the cellular damage produced by the bacteria during the process. This could impose an observational effect which might underestimate the efficiency of the observed recombination and precludes the evaluation of a full SET/RESET cycle by subsequent agroinfiltrations of the actuators in the same leaf. Therefore, we decided to induce SET in a stable PB YFP:LUC plant, to then successfully regenerate *in vitro* new fully switched RL YFP:LUC plants. We monitored the regeneration process with YFP and DNA genotyping finding a number of un-commutated and partially commuted (chimeric) plants. Nevertheless, 30% of the regenerated plants showed complete commutation into RL YFP:LUC RM and constitutive expression YFP, demonstrating that the switch is able to record the SET event keeping its memory. This also indicates that the SET operation can be fully completed in plant systems as it is in bacteria (43). Presumably, such change at the DNA level of somatic T0 cells should be transmitted to the progeny as other authors reported in *Arabidopsis thaliana* for Bxb1 (44) and PhiC31 (45). Subsequently, we induced RESET on the new generated RL YFP:LUC P7 plant, showing increase of Fluc expression and decrease of YFP. This demonstrated that this plant returned to the original PB YFP:LUC, showing a successful SET/RESET cycle over the same RM integrated in the genome. Nevertheless, further optimizations need to be done to overcome the partial efficiency of the RESET, as we still observed presence of the un-commutated RL register.

Finally, we attempted to couple the expression of PhiC31 integrase to an estradiol-inducible system, as a proof of concept of externally inducing the SET operation to control the toggle switch in *benthamiana* hairy roots. Several transgenic root lines showed unwanted activation of the switch due to leaky expression of the integrase. Chemically-inducible systems with low basal expression have been reported in plants (16–19,21); however most of these systems were tested in *Arabidopsis*, and it is possible that chassis-specific conditions (e.g. phytosterol contents) could also determine background expression levels of actuators, which can produce the commutation of the RM. This should be taken into consideration in the design of complex circuits based on PhiC31. Regulation of site-specific DNA excision has been previously reported with the tyrosine recombinase Cre/loxP system in *A. thaliana* using the XVE induction system (46). In this study the authors counter-selected the leaky transformation events by flanking the inducible recombinase and the selection marker with the loxP sites. If leaky, the recombinase would excise this cassette and therefore the transformats would be lost in a selection media. In this work we did not apply negative selection pressure, so all the transformation events were kept. Luckily, we were able to recover one root line with low basal expression but still commutable upon estradiol induction. After activation, low but sustained expression of the Fluc reporter was observed. Similar results were obtained by Hoff et al., 2001 when using heat-shock regulated Cre/loxP (47). They found the system was leaky but still inducible, and that induction worked better in lines with low reporter gene expression like was our case. In order to circumvent this issue, different strategies as the use of degradation tags to regulate the half-life of the integrase or an induction system relying on logic gates to regulate expression of the operators could be implemented. In this regards, a split PhiC31 integrase that can be reconstituted after trans splicing of both components has been developed recently (48).

Summarizing, in this work we created the first modular memory switch for plants making use of the phage PhiC31 integration system. This switch showed to be bistable, reversible and operable, although future efforts will be needed in order to optimize the RESET reaction and avoid unwanted activation. Even though the complexity of big multicellular organisms can be a challenge for extensive characterization of new molecular tools, new advances in the field will allow harnessing their potential. We expect that this system will provide a new tool for PSB that will enable engineering of more complex gene networks and circuits.

## ACKNOWLEDGEMENTS

This work has been carried out at IBMCP-CSIC and funded by Grant BIO201678601-R from Plan Nacional I + D of the Spanish Ministry of Economy and Competitiveness. Bernabé-Orts, JM and Selma, Sara are recipients of an FPI fellowships. We want to thank Dr. Marta Vazquez-Vilar for the advice on the induction systems and Elena Moreno for her support during the *N. benthamiana* hairy roots experiments. The authors declare no conflict of interest.

## SUPPORTING INFORMATION

**Figure S1:** design and cloning details of a custom-made toggle switch.

**Figure S2:** architecture of the register modules (RMs).

**Figure S3:** phenotyping results of the transgenic lines obtained with the transformation with the RMs.

**Figure S4:** influence of the optical density (OD) of the *A. tumefaciens* cultures encoding the SET or RESET operators on the recombination of RMs in *N. benthamiana* stable lines.

**Figure S5:** Quantification of the fluorescence intensity (FI) of the confocal images showed in Fig. 3.

**Figure S6:** Recording the T-DNA expression with the PB and RL register modules.

**Figure S7:** Densitometry of luminescence signal showed in the Fig. 5C image.

**Table S1:** Constructs generated in this study. Sequences are accessible at GB cloning website using the GB database ID.

**Table S2:** oligonucleotides used in this study.

**Table S3:** agroinfiltration cultures and their respective OD600 used in the experiments of Fig. 3 and Fig. S4.

## REFERENCES

1. Vazquez-Vilar, M., Quijano-Rubio, A., Fernandez-Del-Carmen, A., Sarrion-Perdigones, A., Ochoa-Fernandez, R., Ziarsolo, P., Blanca, J., Granell, A. and Orzaez, D. (2017) GB3.0: a platform for plant bio-design that connects functional DNA elements with associated biological data. Nucleic Acids Res., 45, 2196–2209.

2. Sarrion-Perdigones, A., Vazquez-Vilar, M., Palací, J., Castelijns, B., Forment, J., Ziarsolo, P., Blanca, J., Granell, A. and Orzaez, D. (2013) GoldenBraid 2.0: a comprehensive DNA assembly framework for plant synthetic biology. Plant Physiol., 162, 1618–1631.

3. Murray, M.G. and Thompson, W.F. (1980) Rapid isolation of high molecular weight plant DNA. Nucleic Acids Research, 8, 4321–4326.

4. Liu, W. and Neal Stewart, C. (2015) Plant synthetic biology. Trends in Plant Science, 20, 309–317.

5. Baltes, N.J. and Voytas, D.F. (2015) Enabling plant synthetic biology through genome engineering. Trends in biotechnology, 33, 120–131.

6. Ye, X., Al-Babili, S., Kloti, A., Zhang, J., Lucca, P., Beyer, P. and Potrykus, I. (2000) Engineering the provitamin A (beta-carotene) biosynthetic pathway into (carotenoid-free) rice endosperm. Science, 287, 303–305.

7. South, P.F., Cavanagh, A.P., Liu, H.W. and Ort, D.R. (2019) Synthetic glycolate metabolism pathways stimulate crop growth and productivity in the field. Science, 363.

8. Rogers, C. and Oldroyd, G.E.D. (2014) Synthetic biology approaches to engineering the nitrogen symbiosis in cereals. J. Exp. Bot., 65, 1939–1946.

9. Kromdijk, J., Głowacka, K., Leonelli, L., Gabilly, S.T., Iwai, M., Niyogi, K.K. and Long, S.P. (2016) Improving photosynthesis and crop productivity by accelerating recovery from photoprotection. Science, 354, 857–861.

10. Caemmerer, S.v., von Caemmerer, S., Quick, W.P. and Furbank, R.T. (2012) The Development of C4 Rice: Current Progress and Future Challenges. Science, 336, 1671–1672.

11. Xie, M. and Fussenegger, M. (2018) Designing cell function: assembly of synthetic gene circuits for cell biology applications. Nature Reviews Molecular Cell Biology, 19, 507–525.

12. Kassaw, T.K., Donayre-Torres, A.J., Antunes, M.S., Morey, K.J. and Medford, J.I. (2018) Engineering synthetic regulatory circuits in plants. Plant Sci., 273, 13–22.

13. Vazquez-Vilar, M., Orzaez, D. and Patron, N. (2018) DNA assembly standards: Setting the low-level programming code for plant biotechnology. Plant Sci., 273, 33–41.

14. Gardner, T.S., Cantor, C.R. and Collins, J.J. (2000) Construction of a genetic toggle switch in Escherichia coli. Nature, 403, 339–342.

15. Chiu, T.-Y. and Jiang, J.-H.R. (2017) Logic Synthesis of Recombinase-Based Genetic Circuits. Sci. Rep., 7, 12873.

16. Roslan, H.A., Salter, M.G., Wood, C.D., White, M.R., Croft, K.P., Robson, F., Coupland, G., Doonan, J., Laufs, P., Tomsett, A.B. et al. (2001) Characterization of the ethanol-inducible alc gene-expression system in Arabidopsis thaliana. Plant J., 28, 225–235.

17. Zuo, J., Niu, Q.W. and Chua, N.H. (2000) Technical advance: An estrogen receptor-based transactivator XVE mediates highly inducible gene expression in transgenic plants. Plant J., 24, 265–273.

18. Aoyama, T. and Chua, N.H. (1997) A glucocorticoid-mediated transcriptional induction system in transgenic plants. Plant J., 11, 605–612.

19. Mett, V.L., Podivinsky, E., Tennant, A.M., Lochhead, L.P., Jones, W.T. and Reynolds, P.H. (1996) A system for tissue-specific copper-controllable gene expression in transgenic plants: nodule-specific antisense of aspartate aminotransferase-P2. Transgenic Res., 5, 105–113.

20. Saijo, T. and Nagasawa, A. (2014) Development of a tightly regulated and highly responsive copper-inducible gene expression system and its application to control of flowering time. Plant Cell Rep., 33, 47–59.

21. Padidam, M., Gore, M., Lu, D.L. and Smirnova, O. (2003) Chemical-inducible, ecdysone receptor-based gene expression system for plants. Transgenic Res., 12, 101–109.

22. Müller, K., Siegel, D., Rodriguez Jahnke, F., Gerrer, K., Wend, S., Decker, E.L., Reski, R., Weber, W. and Zurbriggen, M.D. (2014) A red light-controlled synthetic gene expression switch for plant systems. Mol. Biosyst., 10, 1679–1688.

23. Inniss, M.C. and Silver, P.A. (2013) Building Synthetic Memory. Current Biology, 23, R812–R816.

24. Roquet, N. and Lu, T.K. (2014) Digital and analog gene circuits for biotechnology. Biotechnol. J., 9, 597–608.

25. Smith, M.C.M., Brown, W.R.A., McEwan, A.R. and Rowley, P.A. (2010) Site-specific recombination by phiC31 integrase and other large serine recombinases. Biochem. Soc. Trans., 38, 388–394.

26. Grindley, N.D.F., Whiteson, K.L. and Rice, P.A. (2006) Mechanisms of site-specific recombination. Annu. Rev. Biochem., 75, 567–605.

27. Friedland, A.E., Lu, T.K., Wang, X., Shi, D., Church, G. and Collins, J.J. (2009) Synthetic gene networks that count. Science, 324, 1199–1202.

28. Bonnet, J., Subsoontorn, P. and Endy, D. (2012) Rewritable digital data storage in live cells via engineered control of recombination directionality. Proc. Natl. Acad. Sci. U. S. A., 109, 8884–8889.

29. Bonnet, J., Yin, P., Ortiz, M.E., Subsoontorn, P. and Endy, D. (2013) Amplifying genetic logic gates. Science, 340, 599–603.

30. Roquet, N., Soleimany, A.P., Ferris, A.C., Aaronson, S. and Lu, T.K. (2016) Synthetic recombinase-based state machines in living cells. Science, 353, aad8559.

31. Siuti, P., Yazbek, J. and Lu, T.K. (2013) Synthetic circuits integrating logic and memory in living cells. Nat. Biotechnol., 31, 448–452.

32. Daniel, R., Rubens, J.R., Sarpeshkar, R. and Lu, T.K. (2013) Synthetic analog computation in living cells. Nature, 497, 619–623.

33. Lyznik, L.A., Gordon-Kamm, W.J. and Tao, Y. (2003) Site-specific recombination for genetic engineering in plants. Plant Cell Reports, 21, 925–932.

34. Wang, Y., Yau, Y.-Y., Perkins-Balding, D. and Thomson, J.G. (2011) Recombinase technology: applications and possibilities. Plant Cell Rep., 30, 267–285.

35. Petolino, J.F. and Kumar, S. (2016) Transgenic trait deployment using designed nucleases. Plant Biotechnol. J., 14, 503–509.

36. Srivastava, V. and Thomson, J. (2016) Gene stacking by recombinases. Plant Biotechnol. J., 14, 471–482.

37. Lutz, K.A., Corneille, S., Azhagiri, A.K., Svab, Z. and Maliga, P. (2004) A novel approach to plastid transformation utilizes the phiC31 phage integrase. Plant J., 37, 906–913.

38. Khaleel, T., Younger, E., McEwan, A.R., Varghese, A.S. and Smith, M.C.M. (2011) A phage protein that binds φC31 integrase to switch its directionality. Mol. Microbiol., 80, 1450–1463.

39. Sheth, R.U. and Wang, H.H. (2018) DNA-based memory devices for recording cellular events. Nat. Rev. Genet., 19, 718–732.

40. Merrick, C.A., Zhao, J. and Rosser, S.J. (2018) Serine Integrases: Advancing Synthetic Biology. ACS Synth. Biol., 7, 299–310.

41. Ghosh, P., Wasil, L.R. and Hatfull, G.F. (2006) Control of phage Bxb1 excision by a novel recombination directionality factor. PLoS Biol., 4, e186.

42. Vazquez-Vilar, M., Bernabé-Orts, J.M., Fernandez-Del-Carmen, A., Ziarsolo, P., Blanca, J., Granell, A. and Orzaez, D. (2016) A modular toolbox for gRNA-Cas9 genome engineering in plants based on the GoldenBraid standard. Plant Methods, 12, 10.

43. Olorunniji, F.J., McPherson, A.L., Rosser, S.J., Smith, M.C.M., Colloms, S.D. and Stark, W.M. (2017) Control of serine integrase recombination directionality by fusion with the directionality factor. Nucleic Acids Res., 45, 8635–8645.

44. Thomson, J.G., Chan, R., Smith, J., Thilmony, R., Yau, Y.-Y., Wang, Y. and Ow, D.W. (2012) The Bxb1 recombination system demonstrates heritable transmission of site-specific excision in Arabidopsis. BMC Biotechnol., 12, 9.

45. Thomson, J.G., Chan, R., Thilmony, R., Yau, Y.-Y. and Ow, D.W. (2010) PhiC31 recombination system demonstrates heritable germinal transmission of site-specific excision from the Arabidopsis genome. BMC Biotechnol., 10, 17.

46. Zuo, J., Niu, Q.W., Møller, S.G. and Chua, N.H. (2001) Chemical-regulated, site-specific DNA excision in transgenic plants. Nat. Biotechnol., 19, 157–161.

47. Hoff, T., Schnorr, K.M. and Mundy, J. (2001) A recombinase-mediated transcriptional induction system in transgenic plants. Plant Mol. Biol., 45, 41–49.

48. Olorunniji, F.J., Lawson-Williams, M., McPherson, A.L., Paget, J.E., Stark, M. and Rosser, S.J. Control of ϕC31 integrase-mediated site-specific recombination by protein trans splicing.

