## Supplementary figures for "A reversible memory switch for plant synthetic biology based on the phage PhiC31 integration system"

**Figure S1:** design and cloning details of a custom-made toggle switch. **(A)** Detailed view of the phytobricks comprising the plant toggle switch. The GOIs are cloned as individual parts in Level 0 using the designated overhangs and then assembled with the PB or RL promoter elements to create a construct for the bistable regulation of each CDS. **(B-D)** Assembly procedure of custom PB and RL register modules using GoldenBraid (GB) cloning system. **(B)** Preparation of the direct and reverse genes of interest (GOIs) for the combination with the PB or RL registers. This involves a BsaI-mediated assembly of Level 1 transcriptional units using Level 0 phytobricks of a promoter, a CDS of the GOI, and a terminator which can be cloned from scratch or reused from the GB's collection. **(C)** Next, these plasmids are used as a PCR-templates to add a couple of overhangs (OH) which define each GOI as reverse or direct CDS and allow its subsequent cloning in a Level 0 plasmid, through a BsmbI-mediated reaction. Standard primers listed in the Table S2 can be used for this amplification. **(D)** Finally, both GOIs cloned as Level 0 parts can be combined with the register PB (GB1494) or the register RL (GB1506) in the Level 1 to conform the PB or RL register modules for the regulated expression of the two GOIs.

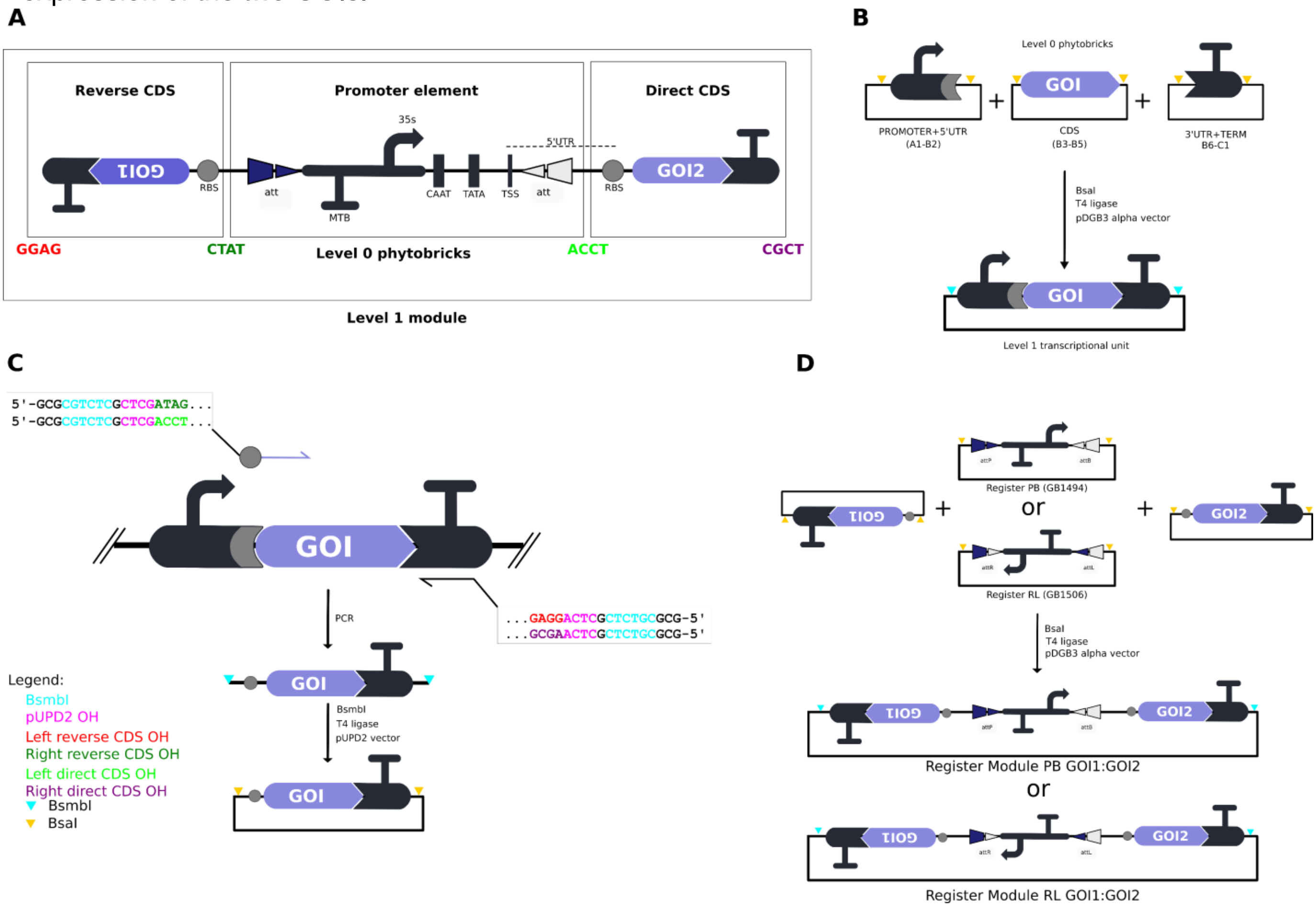

**Figure S2:** architecture of the register modules (RMs) used in the transient expression experiments (A) and for the generation of transgenic lines (B).

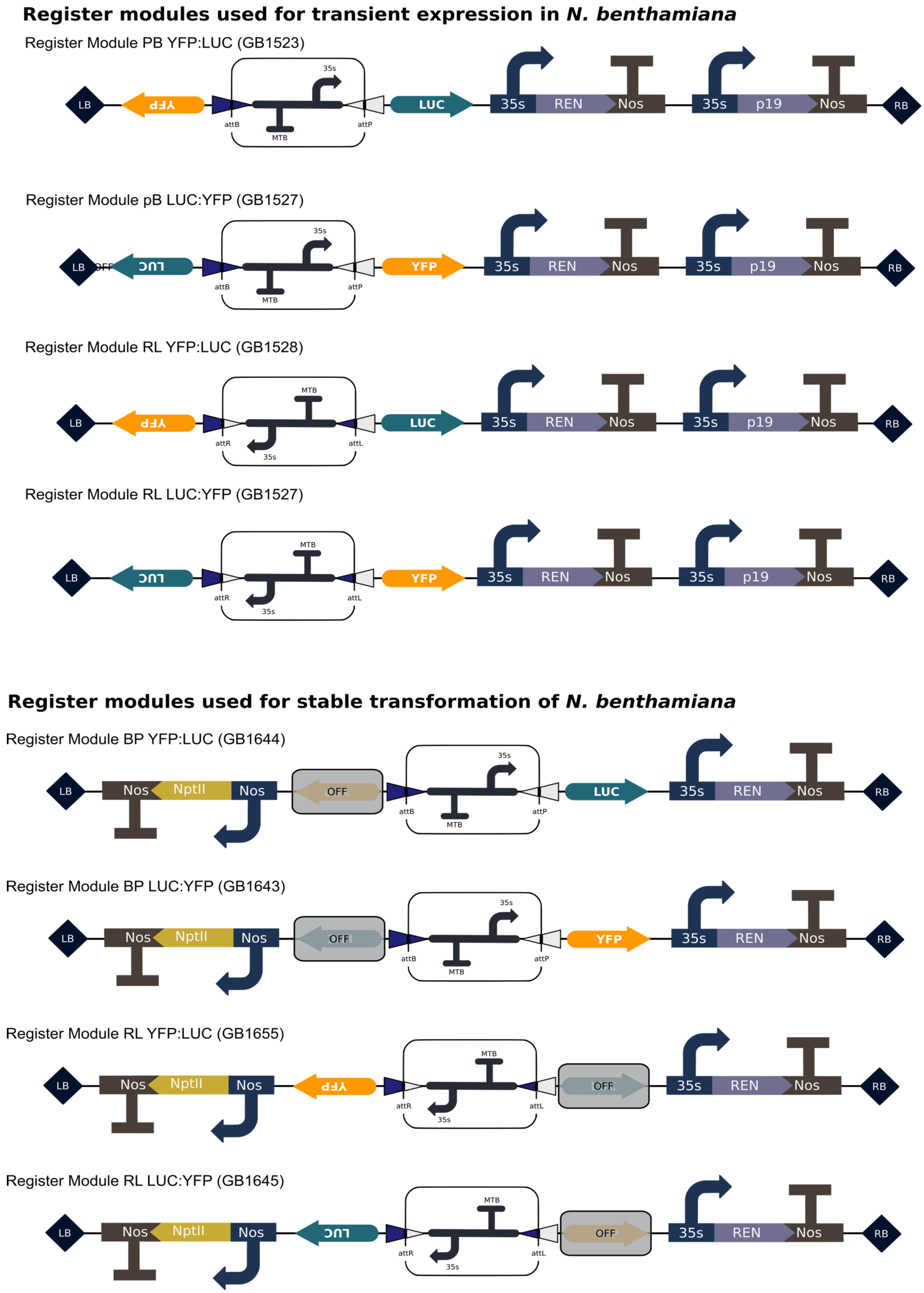

**Figure S3:** phenotyping results of the transgenic lines obtained with the transformation with the RMs. Both, the luminescence (Fluc) and the fluorescence intensity (FI) was measured for each line and are indicated in arbitrary units (a.u.). Marked lines were selected for further experiments in the T1 generation. The bars shows the mean values ± SD of three replicates.

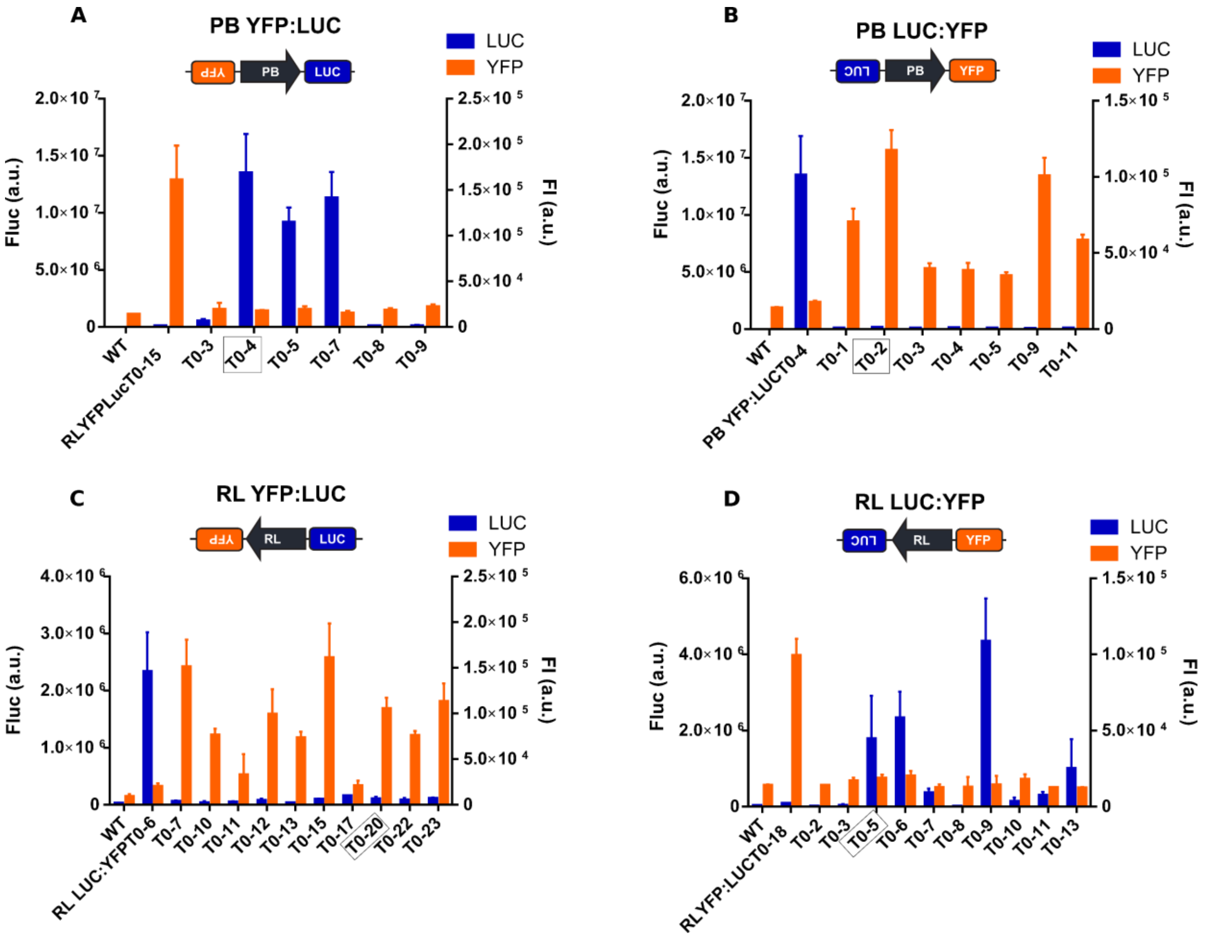

**Figure S4:** influence of the optical density (OD) of the *A. tumefaciens* cultures encoding the SET or RESET operators on the recombination of RMs in *N. benthamiana* stable lines. **A)** Effect of the OD of PhiC31 cultures in the SET process of the PB LUC:YFP T1-2 line. **B)** Effect of the OD of PhiC31 cultures in the SET process of the PB YFP: T1-4 line. **C)** Effect of the ratio of PhiC31:RDF on the reset of the RL LUC:YFP T1-5 line. Ratios are obtained modifying the OD of PhiC31 and RDF cultures. **D)** Effect of the ratio of PhiC31:RDF on the reset of the RL YFP:LUC T1-20 line. Ratios are obtained modifying the OD of PhiC31 and RDF cultures. Bars indicate the fold change between the Fluc/Rluc ratios of treated (PhiC31 or PhiC31-RDF) and the untreated sample (P19), expressed as mean ± SD of three agroinfiltrated leaves.

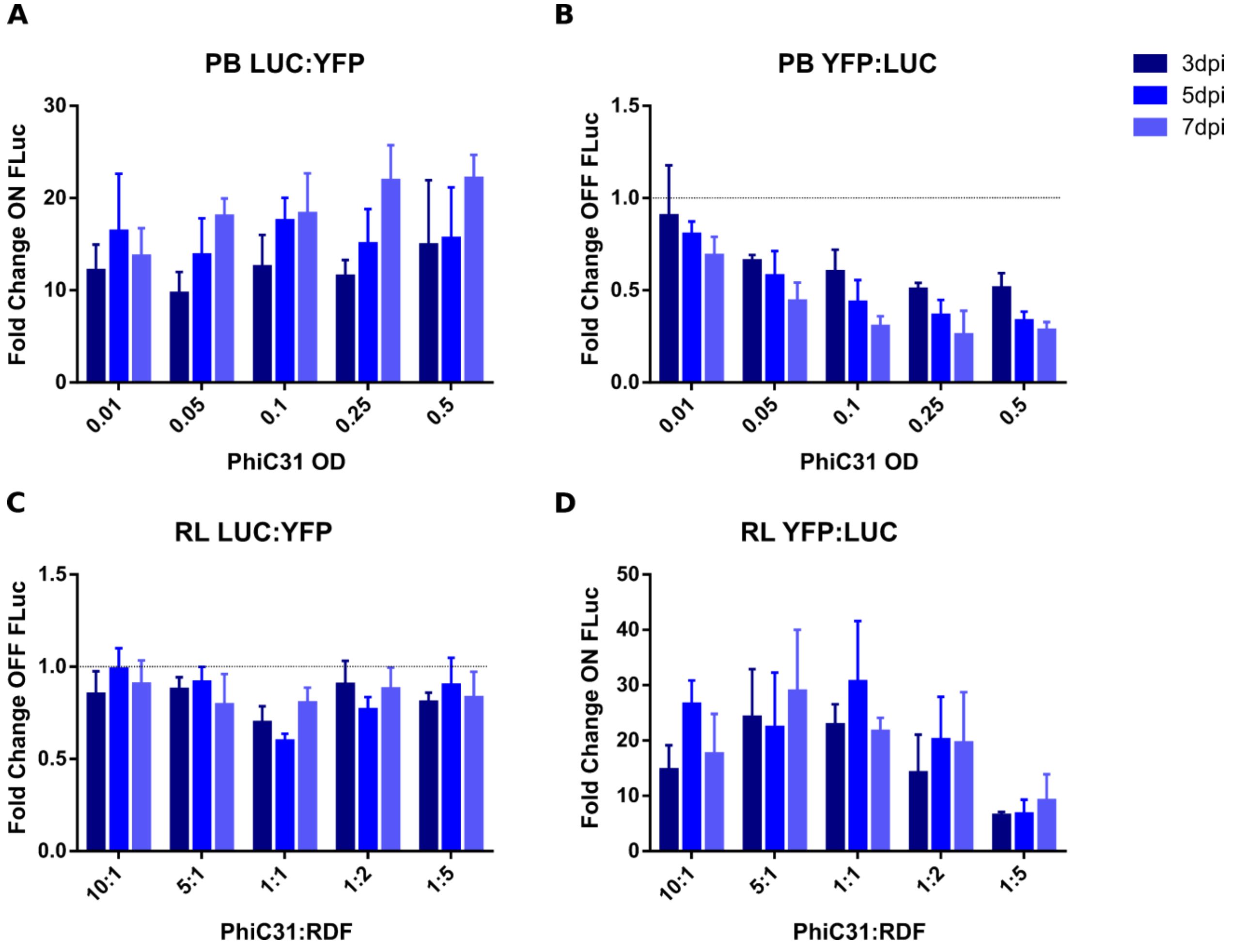

**Figure S5:** Quantification of the fluorescence intensity (FI) of the confocal images showed in Fig. 3. **(A)** FI of the YFP switch on of recombined (+) or not recombined samples (-) for the SET and RESET operations of PB YFP:LUC and RL LUC:YFP register modules. **(B)** Quantification of the YFP switch off of recombined (+) or not recombined samples (-) for the SET and RESET operations of PB LUC:YFP and RL YFP:LUC register modules. Bars shows the mean FI ± SD of nine images.

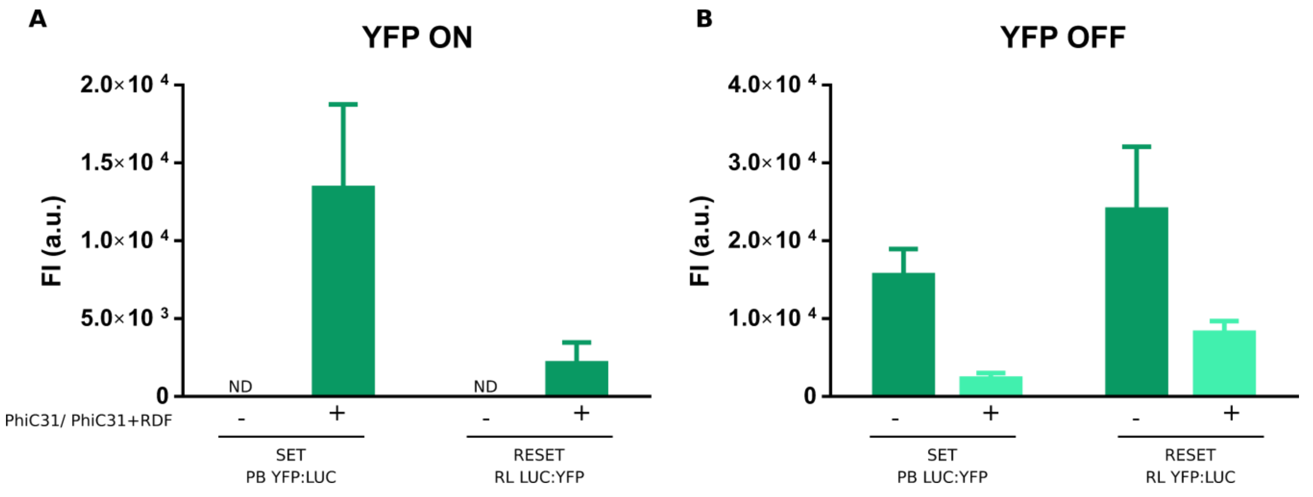

**Figure S6:** Recording the T-DNA expression with the PB and RL register modules. **(A)** Representation of the transformation experiment. Non-fluorescent transgenic RL LUC:YFP and PB YFP:LUC leaf discs were transformed with *A. tumefaciens* carrying a T-DNA with its appropriate actuator (PhiC31+RDF or PhiC31). Leaf discs were cultivated *in vitro* until the callus emerged. If the T-DNA was being expressed, then the presence of the effectors would be registered by the RESET and SET recombination processes which would turn on the YFP expression giving fluorescent callus. **(B)** SET of the PB YFP:LUC and RESET of the RL LUC:YFP generate fluorescent callus visible under the magnification microscope. Scale-bar represents 1 mm. **(C)** Efficiency of the SET and RESET recombination estimated as the number of fluorescent callus respect the total of leaf discs.

**A**

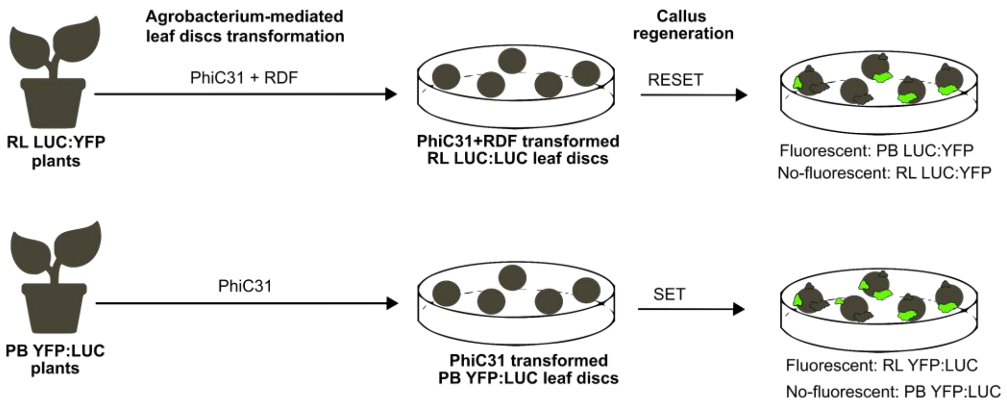


**B**

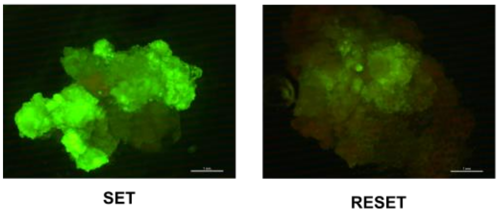


**C**

| | SET | RESET |
|---|---|---|
| Total fluorescent callus | 38 | 16 |
| Total leaf discs | 103 | 110 |
| % Efficiency | **37** | **15** |

**Figure S7:** Densitometry of luminescence signal showed in the Fig. 5C image. **(A)** Quantification of the luminescence signal displayed by estradiol-inducible (EI LUC) and PhiC31-inducible (EI PhiC31) roots incubated for 3 days in estradiol-free (Mock) or estradiol-containing (Estradiol) MS plates. **(B)** Quantification over the same roots 7 days after the estradiol removal. ND means no detected.

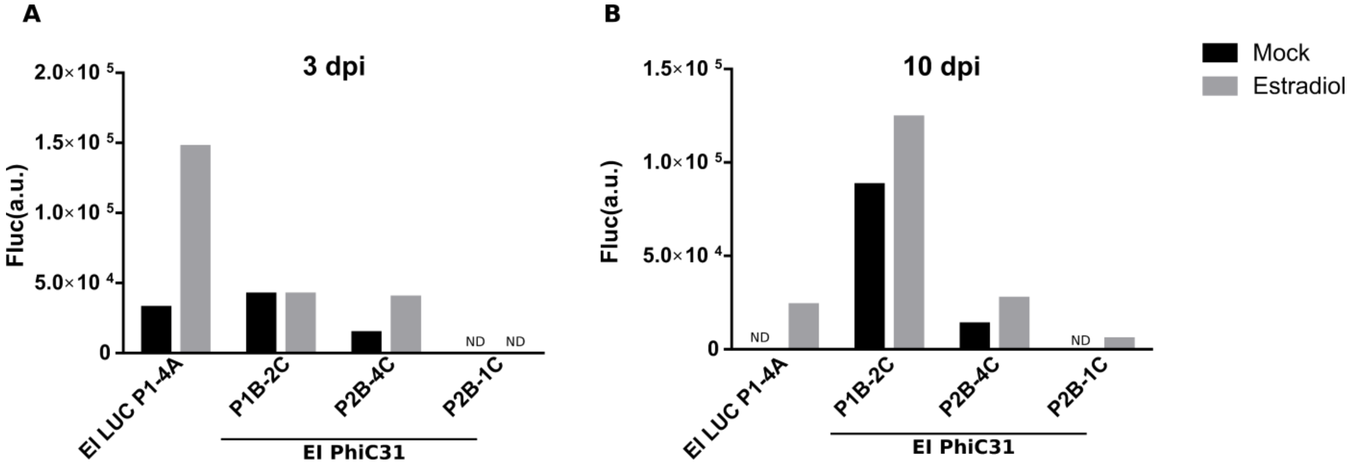
