## Supplementary tables for "A reversible memory switch for plant synthetic biology based on the phage PhiC31 integration system"

**Table S1:** Constructs generated in this study. Sequences are accessible at GB cloning website using the GB database ID.

| Level 0 GB phytobricks | | |
|---|---|---|
| **GB database ID** | **Name** | **Category** |
| 1481 | Assembly Register 1/3 Part- T35S:DsRed:5'UTR | Other |
| 1483 | Assembly Register 3/3 Part - 5'UTR:YFP:Tnos | Other |
| 1494 | Assembly Register 2/3 Part - PhiC31 PB (attP:TMtb:P35S:attB) | Other |
| 1496 | PhiC31 integrase (Plant Codon Optimized) | B3-B4-B5 |
| 1498 | RDF (gp3) | B3-B4-B5 |
| 1499 | Assembly Register 1/3 Part - TNos:Luciferase:5'UTR | Other |
| 1500 | Assembly Register 3/3 Part - 5'UTR:Luc:Tnos | Other |
| 1506 | Assembly Register 2/3 Part - PhiC31 RL (attR:P35S:TMtb:attL) | Other |
| 1507 | Register 1/3 Tnos:YFP:5'UTR | Other |

| Level ≥1 GB phytobricks | | |
|---|---|---|
| **GB database ID** | **Name** | **Category** |
| 460 | P35s:DsRED:T35s-SF | Module |
| 1129 | 35S:ER:LexADBD/Gal4AD:T35S | TU |
| 1130 | OpLexA:mini35S:Luciferase:T35s | TU |
| 1131 | 35s:ER:LexABD/Gal4AD:T35s-OpLexA:mini35s:Luciferase:Tnos | Module |
| 1495 | Register PhiC31 PB (DsRed:YFP) | Other |
| 1497 | P35S:phiC31:T35S | TU |
| 1508 | P35S:RDF:T35S | TU |
| 1510 | Register phiC31 RL (DsRed:YFP) | Other |
| 1513 | Register phiC31 PB (Luc:YFP) | Other |
| 1514 | Register phiC31 RL (Tnos:Luc:attR:P35S:TMtb:attL:YFP:TNos) | Other |

| | | |
|---|---|---|
| 1517 | Register phiC31 PB (YFP:Luc) | Other |
| 1518 | Register phiC31 RL (YFP:Luc) | Other |
| 1523 | Register PB phiC31 Luc:YFP-P35S:Renilla:TNos-P35S:p19:Tnos | Module |
| 1524 | Register RL phiC31 Luc:YFP-P35S:Renilla:TNos-P35S:p19:Tnos | Module |
| 1527 | Register PB phiC31 YFP:Luc-P35S:Renilla:TNos-P35S:p19:Tnos | Module |
| 1528 | Register RL phiC31 YFP:Luc-P35S:Renilla:TNos-P35S:p19:Tnos | Module |
| 1529 | OplexA:mini35S:phiC31:Tnos | TU |
| 1531 | Pnos:phiC31:Tnos | TU |
| 1532 | P35S:ER:lexABD/Gal4AD:T35S - OplexA:mini35S:phiC31:Tnos | Module |
| 1601 | Pnos:ER:LexABD:GAL4AD:Tnos | TU |
| 1643 | Tnos:NptII:Pnos-PB LUC:YFP-P35s:Rluc:T35s | Module |
| 1644 | Tnos:NptII:Pnos-PB YFP:LUC-P35s:Rluc:T35s | Module |
| 1645 | Tnos:NptII:Pnos-RL LUC:YFP-P35s:Rluc:T35s | Module |
| 1655 | Tnos:NptII:Pnos-RL YFP:LUC-P35s:Rluc:T35s | Module |
| 1677 | Pnos:ER:lexABD:GAL4AD:T35s-OplexA:mini35S:phi31:Tnos | Module |
| 2060 | NOS:PhiC31:TNOS - 35s:RDF:T35s | Module |
| 2313 | NOS:ER:LexABD:GAL4AD:T35s - OpLexA:mini35s:PhiC31:TNOS - 35s:DsRED:T35s - SF | Module |
| 2388 | 35s:ER:LexABD:GAL4AD:T35s - OpLexA:mini35s:LUC:Tnos - 35s:DsRED:T35s - SF | Module |

**Table S2**: oligonucleotides used in this study.

| Name | Sequence (5'-3') |
|---|---|
| JO18SEP01 RL YFPLUC F | CAGAGCAGAGATCATGGTGTTAG |
| JO18SEP02 RL YFPLUCR | GCATACGACGATTCTGTGATTTG |
| JO18SEP05 BP YFPLUC F | TTGTGGCTGTTGTAGTTGTACTC |
| JO18SEP06 BP YFPLUCR | ATCATGGTGTTAGCCTTCTATGG |
| JO18SEP03 PhiC31 F | GTTGAATTAGACTGTGGACCGAT |
| JO18SEP04 PhiC31 R | ATCTTGTGCATCGTCTTCATCAT |
| ALF15EN04 | GCGCGTCTCGACGAAAATATAGTTGAAACAGA |
| ALF15EN05 | GCGCGTCTCGTCGTACTAGAGCCAAGCTGATCTC |
| ALF15NOV03 | GCGCGTCTCGCTCGCTATAGTAGTGCCCCAACTGGGGTAACCTTTGAGTTCTCTCAGTTGGGGGCGTA GTCGCAAAAACTATATGCTCT |
| ALF15NOV04 | GCGCGTCTCGCTCAAGGTCGGTGCGGGTGCCAGGGCGTGCCCTTGGGCTCCCCGGGCGCGTACTCC ACTAGTAAATTGTAATGTTGTTTGTTG |
| ALF15DIC06 | GCGCGTCTCGCTCGCTATAGTAGTGCCCCAACTGGG |
| ALF15DIC07 | GCGCGTCTCGCTCAAGGTCGGTGCGGGTGCCA |
| ALF15DIC08 | GCGCGTCTCGCTCGATAGAAACAACATTACAATTTACTATTCTAGTCGA |
| ALF15DIC09 | GCGCGTCTCGCTCGACCTAAACAACATTACAATTTACTATTCTAGTCGA |
| ALF15DIC10 | GCGCGTCTCGCTCAGGAGCGAGTCGGTCCCATT |
| ALF15DIC11 | GCGCGTCTCGCTCAGGAGAGGTCACTGGATTTTGGTTTTAGG |
| ALF15DIC12 | GCGCGTCTCGCTCAAGCGAGGTCACTGGATTTTGGTTTTAGG |
| ALF15DIC13 | GCGCGTCTCGCTCAAGCGCGAGTCGGTCCCATT |

**Table S3:** agroinfiltration cultures and their respective OD600 used in the experiments of Fig. 3 and Fig. S4.

| Optimization of PhiC31 Culture | Optimization of PhiC31 OD | Optimization of PhiC31+RDF Culture | Optimization of PhiC31+RDF OD | PB register kinetics Culture | PB register kinetics OD | RL register kinetics Culture | RL register kinetics OD |
|---|---|---|---|---|---|---|---|
| PhiC31 | 0.01 | PhiC31 | 0.1 | PhiC31 | 0.1 | PhiC31 | 0.1 |
| | 0.05 | RDF | 0.01 | NA | NA | RDF | 0.1 |
| | 0.1 | | 0.05 | NA | NA | | |
| | 0.25 | | 0.1 | NA | NA | | |
| | 0.5 | | 0.2 | NA | NA | | |
| | | | 0.5 | NA | NA | | |
| P19 | 0.1 | P19 | 0.1 | 2 vol. P19 | 0.1 | P19 | 0.1 |
